# Extrasynaptic signaling enables an asymmetric juvenile motor circuit to produce a symmetric undulation

**DOI:** 10.1101/2021.09.21.461278

**Authors:** Yangning Lu, Tosif Ahamed, Ben Mulcahy, Jun Meng, Daniel Witvliet, Sihui Asuka Guan, Douglas Holmyard, Wesley Hung, Quan Wen, Andrew D Chisholm, Aravinthan DT Samuel, Mei Zhen

## Abstract

In many animals, there is a direct correspondence between the motor patterns that drive locomotion and the motor neuron innervation onto the muscle groups. For example, the adult *C. elegans* moves with symmetric and alternating dorsal-ventral bending waves arising from symmetric motor neuron input onto the dorsal and ventral muscles. In contrast to the adult, the *C. elegans* motor circuit at the juvenile larval stage has asymmetric wiring between motor neurons and muscles, but still generates adult-like bending waves with dorsal-ventral symmetry. We show that in the juvenile circuit, wiring between excitatory and inhibitory motor neurons coordinates the contraction of dorsal muscles with relaxation of ventral muscles, producing dorsal bends. However, ventral bending is not driven by analogous wiring. Instead, ventral muscles are excited uniformly by premotor interneurons through extrasynaptic signaling. Ventral bends occur in anti-phasic entrainment to activity of the same motor neurons that drive dorsal bends. During maturation, the juvenile motor circuit is replaced by two motor subcircuits that separately drive dorsal and ventral bending. Modeling reveals that the juvenile’s immature motor circuit is an adequate solution to generate adult-like dorsal-ventral bending before the animal matures. Developmental rewiring between functionally degenerate circuit solutions, that both generate symmetric bending patterns, minimizes behavioral disruption across maturation.

**Highlights:**

- *C. elegans* larvae generate symmetric motor pattern with an asymmetrically wired motor circuit.
- Synaptic wiring between excitatory and inhibitory motor neurons drives dorsal bending.
- Extrasynaptic excitation by premotor interneurons entrains ventral muscles for anti-phasic ventral bending.
- A developmental strategy to enable mature motor pattern before the circuit structurally matures.

## Introduction

Many animals generate alternating motor patterns that are thought to arise from symmetric input from motor neurons to muscle groups that execute alternating movements. In vertebrates, the hind-limb’s movement is driven by rhythmogenic spinal interneuron circuits^1,2^. Structurally symmetric interneuron circuits coordinate the contraction and relaxation of muscle groups in either the left or right limb. Phase relations between the interneuron circuit for the left and right limbs produce alternating gaits^3–5^. In the leech, separate excitatory and inhibitory motor neurons innervate left-right side of the muscle groups during swimming^6^. In two nudibranch molluscs, a bilaterally pair of rhythmogenic interneuron circuits drives the left or right flexions, respectively, resulting in an alternating swimming pattern^7^.

The adult *C. elegans* motor system consists of two homologous motor neuron networks^8,9^ (illustrated in Figure 1B, right panel). In this circuit, dorsal and ventral muscles form neuromuscular junctions (NMJs) with two sets of cholinergic excitatory motor neurons (eMNs) that contract muscles, as well as with two sets of GABAergic inhibitory motor neurons (iMNs) that relax muscles^10–12^. NMJs from eMNs are also dyadic synapses to the iMNs that form NMJs with muscles on the opposite side, thus coordinating muscle contraction and relaxation.

**Figure 1.**
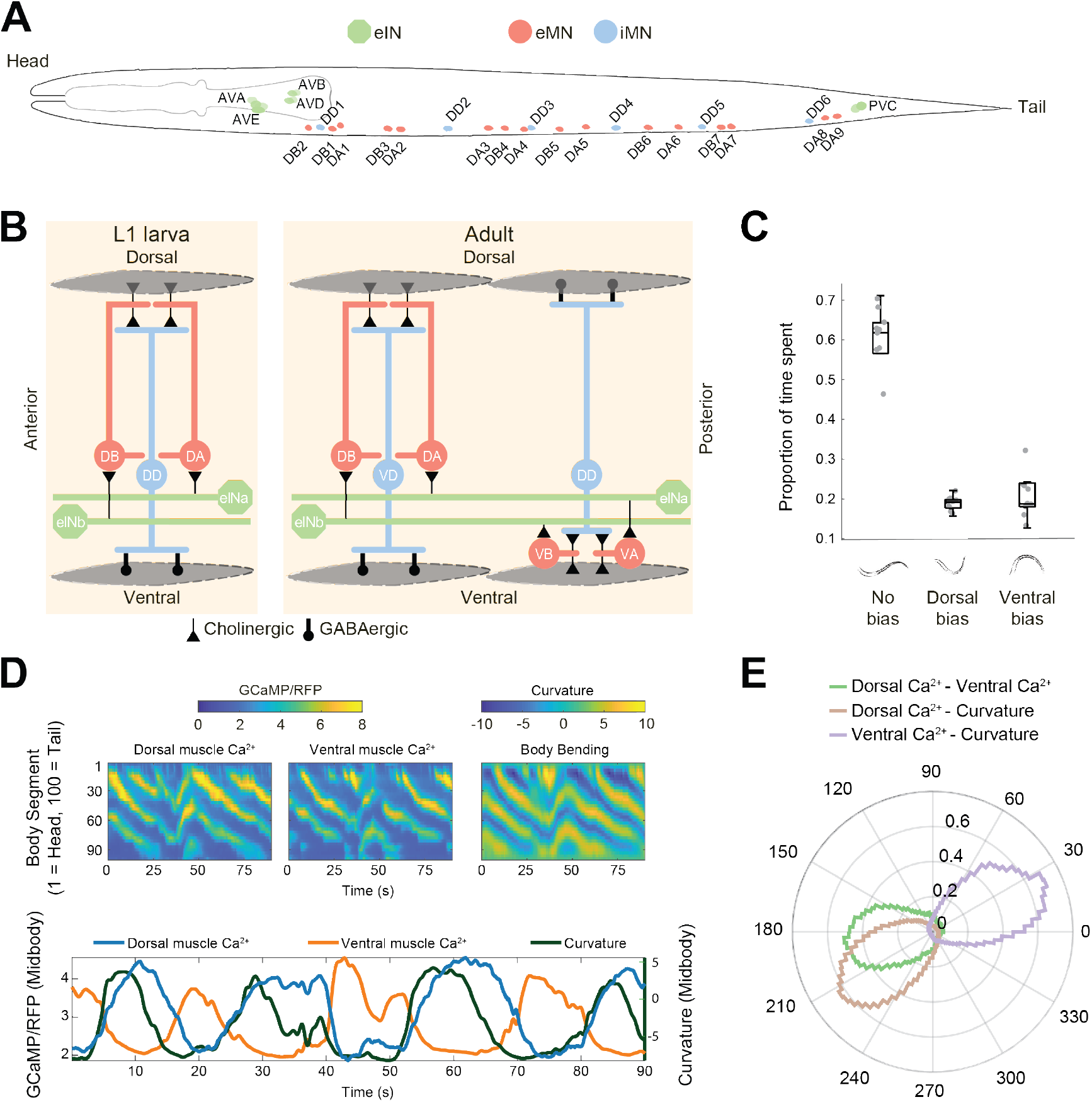
L1 larvae exhibit asymmetry in motor circuit wiring but symmetry in their motor pattern and muscle activity. (A) Schematic of the motor circuit for body movement in a newborn *C. elegans* L1 larva. Circles denote the position of neuron soma, color-coded by class. eINs, colored in green, include a pair of AVA, AVB, AVD, AVE, and PVC premotor interneurons. eMNs, colored in red, include DA and DB motor neurons. There are 9 DA motor neurons and 7 DB motor neurons. iMNs, colored in blue, include 6 DD motor neurons. (B) (Left) schematic of L1 larva’s motor circuit synaptic wiring, deduced by serial section EM reconstruction. (Right) schematic of adult’s motor circuit synaptic wiring^9^. Diamond shapes denote muscles. Cholinergic synapses are marked as triangles. GABAergic synapses are marked as circles. (C) L1 larvae swim without a dorsal-ventral bias most of the time and spend roughly equal time in postures with a dorsal or ventral bias. n = 10 larvae. (D) Calcium dynamics of dorsal and ventral muscles in a crawling L1 larva. Colormap represent the GCaMP3/RFP ratio in muscles (top left panels) and the body curvature (top right panel), respectively. (bottom panel) Time series of curvature (right Y-axis), dorsal and ventral muscle activity (left Y-axis) at mid-body (segment 50). (E) Polar histograms of the phase differences between curvature, dorsal muscle activity, and ventral muscle activity along the body (segments 33-95). n = 12 larvae.

One motor subcircuit consists of repeated modules of cholinergic eMNs (DA and DB) and GABAergic iMNs (VD). They coordinate simultaneous dorsal muscle contraction and ventral muscle relaxation in each body segment, producing dorsal bends. A complementary motor subcircuit produces ventral bends, with symmetric wiring of a homologous set of eMNs (VA and VB) and iMNs (DD)^9^ (illustrated in Figure 1B, right panel). Integrating the role of oscillators and proprioceptors, eMNs underlie rhythmic and coordinated body bending^13–18^. Alternating activity of these two subcircuits in each body segment gives rise to alternating dorsal and ventral bending during adult locomotion^19^.

However, the newly born larva (L1) lacks this symmetry in motor neuron output (Figures 1A and B, left panel). For the first few hours after birth, an L1 larva has only the DA, DB, and DD motor neurons^20^. A partial serial section electron microscopy (ssEM) reconstruction of the L1 larva^8^ suggests that its motor neuron wiring is different from that in the adult^9^. In examined larva body segment, eMNs (DA and DB) make NMJs exclusively to dorsal muscles and iMNs (DD) make NMJs exclusively to ventral muscles. NMJs from eMNs are also dyadic synapses to the iMNs. Hence the entire L1 motor neuron wiring resembles adult’s motor neuron subcircuit for dorsal bending (Figure 1B).

Despite lacking a homologous motor circuitry to drive ventral bends, the L1 larva crawls with an adult-like undulation. Here, we sought mechanisms by which the L1’s structurally asymmetric motor circuit produces its motor pattern with dorsal-ventral symmetry. First, we used serial section EM to fully reconstruct the connectivity between premotor interneurons, motor neurons, and muscle cells in the L1 motor circuit. We then used functional imaging, optogenetic perturbations, as well as cell and synapse ablation to assess the role of each circuit component in generating dorsal-ventral bends. Finally, we combined modeling and behavioral simulations to develop an integrated understanding of underlying mechanisms.

We found that the L1 motor circuit functions through both synaptic and extrasynaptic connections. Its motor neurons form a self-regulated circuit to generate and exit dorsal bends. Ventral bends are produced by anti-phasic entrainment: extrasynaptic transmission from cholinergic premotor interneurons (eINs) excites ventral muscles uniformly along the body, which allows the same motor neurons for dorsal bending to produce complementary ventral bends.

We present an example of circuit degeneracy^21–23^ in an alternative solution for generating a symmetric gait from an asymmetrically wired motor circuit. With extrasynaptic transmission and neurons adopting additional roles, an immature motor circuit can generate a mature motor pattern. These are adaptive strategies by which an animal maintains behavioral output without functional disruption across maturation, despite substantial structural changes.

## Results

### The L1 motor neurons have asymmetric input to dorsal and ventral body wall muscles

*C. elegans* is born with a fraction of motor neurons of the adult motor circuit. Beginning at the mid-L1 larva stage, post-embryonic neurogenesis gives rise to new motor neurons^24^. Because all motor neurons that innervate ventral muscles in adults are born post-embryonically^9^, the L1 larva motor neurons must have a distinct, and potentially asymmetrically wired configuration.

We used serial section EM to fully reconstruct the motor circuit of multiple L1 larvae, from premotor interneurons to motor neurons to muscles (^25^ and this study). In an early L1 larva (Methods; Figures 1 and S1), for motor neurons, we confirmed results from a previous partial reconstruction^8^: inputs to dorsal and ventral muscles are asymmetric (Figure 1B, left panel). All cholinergic eMNs (DA and DB) make NMJs only to dorsal muscles. All GABAergic iMNs (DD) make NMJs only to ventral muscles. Most NMJs from the eMNs are dyadic, innervating dorsal muscles as well as the iMNs that project to the opposite side (examples in Figure S1). Multiple eMNs and iMNs (Figure 1A) form a chain of similarly wired modules along the body (Figure 1B, left panel).

In the adult motor circuit, two different groups of premotor interneurons (eINs) innervate the A- and B-class eMNs to regulate directional movements^19,26^. Our EM reconstruction revealed that this wiring pattern is already present in the newborn L1 larva (Figures 1 and S1;^25^). The DA subclass of eMNs are postsynaptic to cholinergic eINs that regulate backward locomotion (AVA, AVE, and AVD); the DB subclass of eMNs are postsynaptic to eINs that promote forward locomotion (AVB and PVC). Thus the key differences in structural wiring between the L1 and adult motor circuits are the motor neurons and their connections to muscles (Figure 1B, right panel).

### L1 larvae generate alternating dorsal-ventral body bends and muscle activities

Despite dorsal and ventral muscles receiving asymmetric inputs from motor neurons, *C. elegans*’s alternating dorsal-ventral bending pattern is established at birth. When swimming, L1 larvae exhibited sinusoidal bending waves and full-body coils without dorsal or ventral bias (Figure 1C; Videos S1A). When crawling, L1 larvae propagated alternating, dorsal-ventral bending waves along the body (Figures 1D and 1E; Video S1B).

The calcium dynamics of body wall muscles correlated with this bending pattern. In slowly crawling L1 larvae (Methods; Video S1C), calcium waves propagated along dorsal and ventral muscles and tracked curvature changes (Figures 1D and 1E). Consistently, dorsal-ventral alternation in body bending correlated with out-of-phase activation of the corresponding muscles (Figures 1D and 1E). Overall, dorsal and ventral muscle activity level is balanced.

### Cholinergic excitatory motor neurons drive dorsal muscle contraction

Wiring asymmetry of the L1 motor circuit needs to be reconciled with bending symmetry of the L1 larva. One hypothesis is that cholinergic and GABAergic synapses may have mixed signs (inhibitory or excitatory) in L1, compensating for the wiring asymmetry. To determine the functional relationship between cholinergic motor neurons and bending, we measured calcium dynamics of eMNs in moving L1 larvae.

We found that when L1 larvae moved backwards, DA motor neurons were activated in retrograde sequence as bending traveled from tail to head (Figure 2A; Video S2A). Calcium dynamics at each DA soma correlated with dorsal bending, exhibiting the shortest time lag with bending of the anterior body segment (Figure 2A, lower right panel). This position-dependent correlation is consistent with the anatomical wiring: DA axons project anteriorly, making NMJs to dorsal muscles anterior to their somas (Figure 1B, left panel).

**Figure 2.**
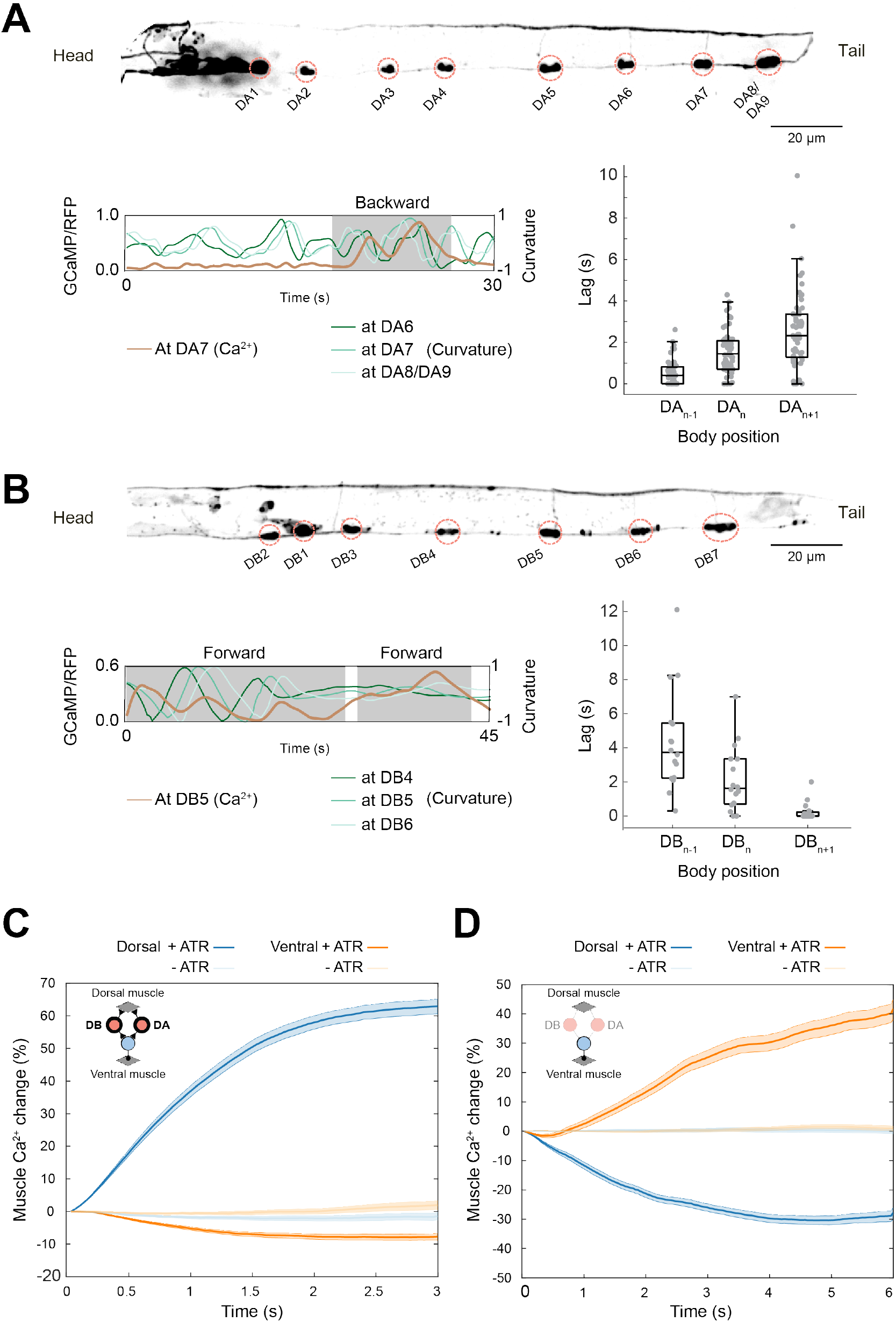
Cholinergic eMNs underlie dorsal muscle contraction. (A) Top, GCaMP6s expression in the DA subclass eMNs in an L1 larva. Bottom left, an example trace of calcium activity for DA7 motor neuron (left Y-axis), overlaid with curvatures at different body segments (right Y-axis). Shaded area denotes backward movement. Bottom right, shortest time lag between calcium activity and curvature changes at regions anterior to (N-1), at (N), and posterior to (N+1) the neuron’s soma (DA_n_), during periods of backward movement. (B) Left panel: Top: GCaMP6s expression in the DB subclass eMNs. Bottom left: an example calcium trace for DB5 motor neuron (left Y-axis), overlaid with curvatures at different body segments (right Y-axis). Bottom right: shortest time lag between calcium activity and curvature changes at body segments anterior to (N-1), at (N), and posterior to (N+1) the neuron’s soma (DB_n_), during periods of forward movement. (C) Simultaneous activation of all eMNs by Chrimson and calcium imaging of body wall muscles in L1 larvae. Y-axis plots percentage changes of the muscle activity from *t* = 0. Control group (-All-Trans Retinal (ATR)): 21 stimulation epochs from 7 larvae. Experimental group: (+ ATR): 33 stimulation epochs from 12 larvae. (D) Simultaneous inactivation of eMNs by GtACR2 and calcium imaging of body wall muscles in L1 larvae. Control group (-ATR): 39 stimulation epochs from 8 larvae. Experimental group (+ ATR): 35 stimulation epochs from 12 larvae. Lines and shades denote median and 95% confidence intervals, respectively for (C) and (D).

When L1 larvae moved forward, DB motor neurons were activated in an anterograde sequence as bending traveled from head to tail (Figure 2B; Video S2B). Calcium dynamics at each DB soma correlated with dorsal bending, exhibiting the shortest time lag with bending of the posterior body segment (Figure 2B, lower right panel), consistent with the posterior projection of DB axons (Figure 1B, left panel). Both results establish positive correlations between eMNs and dorsal bending.

To determine whether eMNs directly activate dorsal muscles, we imaged muscle calcium dynamics while simultaneously manipulating motor neuron activity (Methods). We found that activation of all eMNs by Chrimson^27^ increased the activity of dorsal muscles, but not ventral muscles (Figure 2C). Similarly, inhibition of eMNs by GtACR2^28^ lowered the activity of dorsal muscles, but led to an activity increase in ventral muscles (Figure 2D). Cholinergic inputs from eMNs are thus excitatory specifically to dorsal muscles.

### GABAergic inhibitory motor neurons promote ventral and dorsal muscle relaxation

GABAergic motor neurons (DD) make NMJs exclusively to ventral body wall muscles. During early postnatal development, GABAergic synapses from rodent interneurons are not inhibitory but excitatory^29^. In *C. elegans*, whether GABAergic signaling promotes muscle relaxation in L1 larvae^30^, as in adults^31^, was unclear^32^.

To assess how DD motor neurons regulate muscle activity, we measured their dynamics in crawling L1 larvae. Each DD motor neuron makes a cluster of NMJs that can be recorded as a single region of ROI (Figure 3A, NMJ). Whether these animals moved backward or forward, activity rise of DD NMJs correlated with ventral relaxation (Figure 3B; Video S3A), and the calcium rise preceded decreased curvature at each ROI (Figure 3A, bottom panel and Figure 3B, right panel).

**Figure 3.**
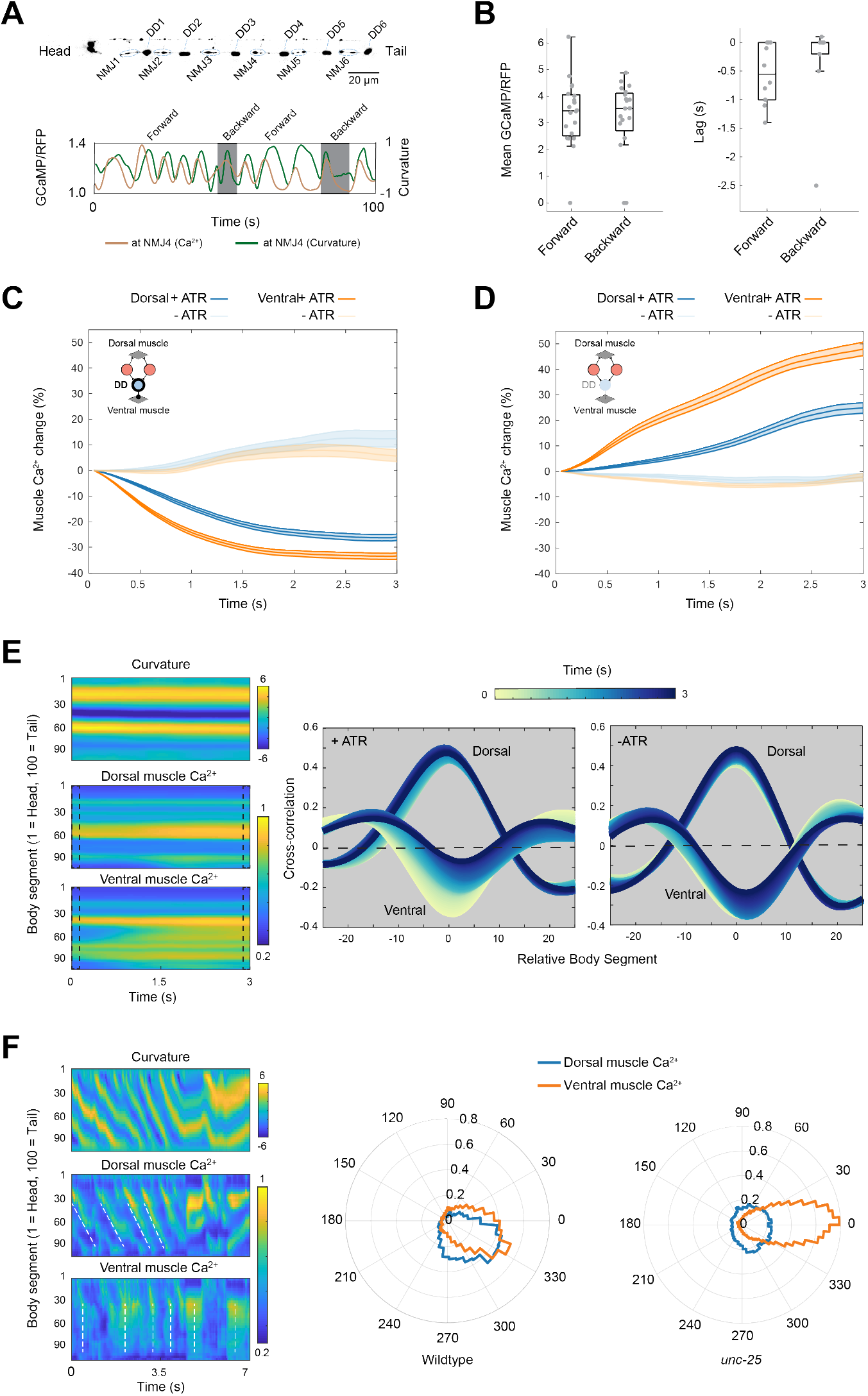
GABAergic iMNs underlie ventral and dorsal muscle relaxation. (A) (Top) GCaMP6s expression in the DD motor neurons in an L1 larva. Each DD makes a tight cluster of NMJs; NMJn is from DDn. (Bottom) An example calcium trace of NMJ4 (left Y-axis), overlaid with body curvature at the same segment (right Y-axis). Shaded areas denote periods of backward movements. (B) (Left) Mean calcium activity of DD NMJs during forward and backward movement. n = 17 NMJs from 12 larvae; (Right) Shortest time lags between calcium activity and curvature changes during forward and backward movements. (C) Simultaneous activation of all iMNs (DD) by Chrimson and muscle calcium imaging in L1 larvae. Y-axis plots percentage changes of the muscle activity from *t* = 0. Control group (-ATR): n = 13 stimulation epochs from 7 larvae; Experimental group (+ ATR): n = 18 stimulation epochs from 10 larvae. (D) Simultaneous inhibition of all iMN (DD) by Archaerhodopsin and calcium imaging of body wall muscles in L1 larvae. Control group (-ATR): n = 10 stimulation epochs from 5 larvae; Experimental group (+ ATR): n = 19 stimulation epochs from 10 larvae. Lines and shades represent median and 95% confidence interval, respectively for (C) and (D). (E) (Left) Example heatmap of curvature, and dorsal and ventral muscle calcium of an L1 larva during inactivation of iMNs (DD) by Archaerhodopsin. Dashed black boxes highlight the differences in muscle calcium at beginning and end of the recording. At the beginning, ventral muscle calcium is in accordance with the curvature. However as the iMNs (DD) are inhibited, this relationship weakens, seen in the dotted box towards the end of the recording. In contrast, dorsal muscle calcium continues a correlated pattern with curvature through the recording. (Right) Spatial cross-correlation between the muscle calcium and curvature along the body (segment 33-95) during the course of iMN inactivation. Color encodes the time-points at which the cross-correlation was calculated. Only for ventral muscles in the presence of ATR, the correlation decreased towards zero over the course of the recording. Quantified from the same data as in (D). (F) (Left) Example heatmap of curvature and dorsal and ventral muscle calcium in a crawling mutant larva that cannot synthesize GABA (*unc-25*). Dashed white lines denote propagating calcium waves in dorsal muscles, and non-propagating waves in ventral muscles. (Right) Polar histograms of muscle activity phase lags between segments along the body in wildtype and *unc-25* mutant L1 larvae. The body (segments 33-95) is binned into 6 equally spaced sections. Phase differences are calculated between subsequent sections separately for dorsal and ventral muscles. Phase difference between ventral muscle activity between subsequent sections shows a peak at 0 in the histogram. n = 10 larvae.

To determine whether DD motor neurons inhibit ventral muscles, we imaged muscle calcium dynamics while manipulating DD activity. As expected from inhibitory NMJs, activation of DD by Chrimson strongly reduced ventral muscle activity (Figure 3C). Unexpectedly, their activation also led to a smaller reduction of dorsal muscle activity (Figure 3C), even though DD motor neurons do not make NMJs to dorsal muscles (Figure 1B, left panel).

Inhibition of DD motor neurons by Archaerhodopsin^33^ had an opposite effect: a strong increase in ventral muscle activity, accompanied by a smaller increase in dorsal muscle activity (Figure 3D; Video S3B). More intriguingly, activity increase of dorsal and ventral muscles exhibited spatial differences: while dorsal activity continued to track curvature, ventral activity increase was uniform along the body (Figure 3E; Video S3B). We quantified this effect by comparing the correlation between curvature and calcium activity in the dorsal and ventral body: a uniform activity increase along the body regardless of curvature led to decreasing correlation with curvature over time, and this decrease was only observed for ventral muscles and required iMN inactivation (Figure 3E, right panels).

We observed a similar spatial difference between dorsal and ventral muscle activity when we genetically silenced DD motor neurons (Figure 3F). In crawling mutant larvae that do not synthesize GABA (*GAD/unc-25*), dorsal muscles exhibited calcium dynamics that correlated with propagating bending waves, whereas calcium signals of ventral muscles did not propagate (Figure 3F; Video S3C; Figure S2). The non-propagating ventral muscle calcium signal was reminiscent of the calcium activity distribution regardless of curvature (Figure 3E). Standing calcium patterns similarly disrupt the correlation with curvature (Figure 3F, right panels).

Therefore, DD motor neurons are inhibitory at birth. These iMNs inhibit ventral muscles with synapses and inhibit dorsal muscles without direct synaptic input. Dorsal and ventral muscles exhibit structural, temporal, and spatial differences in their relationships with iMNs, implying distinct inhibitory mechanisms.

### Synaptic wiring between excitatory and inhibitory motor neurons form a circuit for dorsal bending

Most NMJs from eMNs to dorsal muscles are dyadic synapses, juxtaposing dendrites of iMNs that inhibit the opposing ventral muscles (Figure S1). Thus, activation of dorsal muscles should lead to contralateral ventral muscle inhibition.

Consistent with this hypothesis, activation of all eMNs by Chrimson increased the iMN activity (Figure 4A; Videos S4A and S4B). Inhibition of all eMNs by GtACR2 decreased the iMN activity (Figure 4A; Videos S4C and S4D). Furthermore, ablation of the DB subclass eMNs by miniSOG^34^ preferentially reduced DD’s activity when animals attempted to move forward (Figure S3A), whereas ablation of the DA subclass eMNs reduced DD’s activity when animals attempted to move backward (Figure S3B).

**Figure 4.**
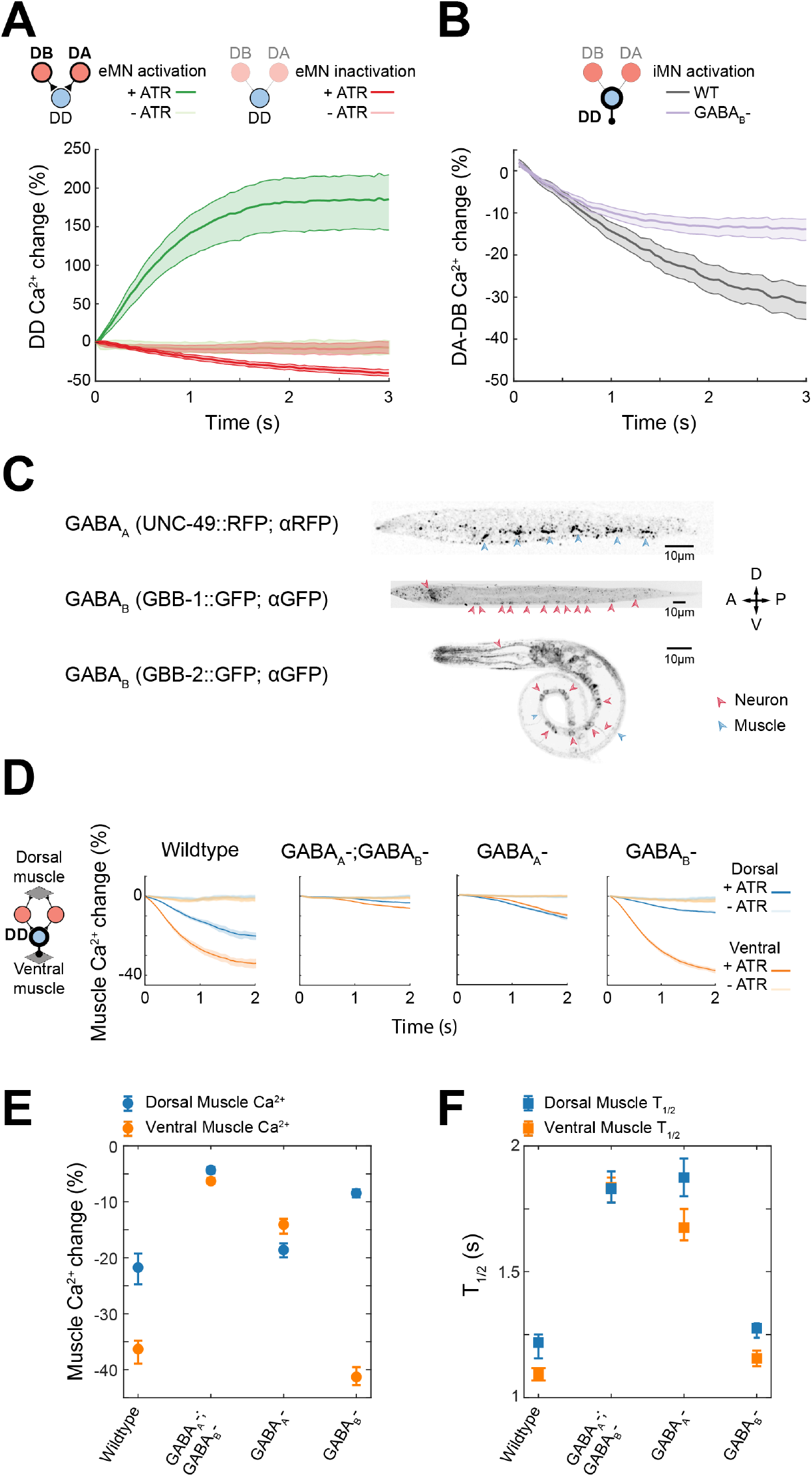
iMNs are activated by eMNs and inhibit muscles through ionotropic and metabotropic GABA signaling. (A) Optogenetic activation and inactivation of eMNs leads to corresponding changes in iMN activity. (Inset) Schematic depicting the wiring from eMNs to iMNs. (Green) Activation of eMNs by Chrimson elevates iMN activity. - ATR group: n = 20 stimulation epochs from 5 larvae; + ATR group: n = 23 stimulation epochs from 9 larvae. (Red) Inhibition of eMNs by Archaerhodopsin decreases DD activity. - ATR group: n = 25 stimulation epochs from 6 larvae; + ATR group: n = 21 stimulation epochs from 9 larvae. (B) Optogenetic activation of iMNs leads to decrease of eMN activity. (Inset) Schematic depicting the lack of synaptic wiring from iMNs to eMNs. Colored lines denoted the response in wildtype and *gbb-2* mutant larvae in the presence of ATR. Wildtype larvae: 28 stimulation epochs from 4 larvae; *gbb-2* mutant larvae: 137 stimulation epochs from 6 larvae. Lines and shades denote median and 95% confidence intervals, respectively in (A) and (B). (C) Expression patterns of endogenously tagged GABA_A_::RFP receptors and GFP tagged fosmid reporters for two subunits of a heterologous GABA_B_ receptor complex, GBB-1 and GBB-2. (D) (Inset) Schematic depicting simultaneous optogenetic activation of all iMNs and muscle calcium imaging. Y-axis plots percentage changes of the ventral and dorsal muscle activity from *t* = 0. iMN activation reduced both dorsal and ventral muscle activities in wildtype L1 larvae. The reduction was differentially attenuated in ionotropic GABA_A_ and metabotropic GABA_B_ double or single mutant larvae. For GABA_A_ and GABA_B_ double mutants GABA_A_-; GABA_B_- (*unc-49(e407); gbb-2(tm1165)*): Control group (-ATR): 46 stimulation epochs from 12 larvae; Experimental group (+ ATR): 76 stimulation epochs from 19 larvae. For GABA_A_ mutants GABA_A_- (*unc-49(e407)*: Control group (-ATR): 37 stimulation epochs from 22 larvae; Experimental group (+ ATR): 51 stimulation epochs from 20 larvae. For GABA_B_ mutants GABA_B_- (*gbb-2(tm1165)*): Control group (-ATR): 18 stimulation epochs from 6 larvae; Experimental group (+ ATR): 39 stimulation epochs from 7 larvae. Lines and shades denote median and 95% confidence intervals, respectively. (E) Maximal activity changes in dorsal and ventral muscles in respective genotypes shown in (D). (F) Kinetics measured through the half-time of dorsal and ventral muscle activity changes in respective genotypes shown in (D).

We have shown that NMJs to dorsal muscles and ventral muscles drive contraction and relaxation, respectively (Figures 2 and 3). Dyadic NMJs from eMNs to iMNs promote relaxation of juxtaposed ventral muscles when dorsal muscles contract. This coordination therefore generates dorsal bending.

### Extrasynaptic GABA signaling plays a dominant role in dorsal muscle inhibition

An exit from dorsal bending requires dorsal muscles to transit from contraction to relaxation. We have shown that iMN activation leads to inhibition of both ventral and dorsal muscle activities (Figure 3). However, iMNs only make inhibitory NMJs to ventral muscles, not to dorsal muscles, or to eMNs that activate dorsal muscles (Figures 1B, left panel, and S1). This raises the possibility of iMNs inhibit ventral muscle synaptically, while inhibiting dorsal muscles by extrasynaptic signaling.

Previous studies have established that synaptic GABA inhibition involves the ionotropic GABA_A_ receptors, and extra-synaptic GABA inhibition involves the metabotropic GABA_B_ receptors^35–37^. Consistent with previous reports^38,39^, in newborn L1 larvae, the GABA_A_ receptor (UNC-49) is present only in ventral muscles, and absent from the dorsal muscles (Figure 4C, top panel). Along the ventral body, endogenously tagged UNC-49::RFP formed six clusters (Figure 4C), reminiscent of the NMJ clusters by individual DD motor neurons (Figure 3A), as we observed by the EM reconstruction. Further supporting the notion that iMNs inhibit ventral muscles by synaptic transmission, in the absence of GABA_A_ receptors (*unc-49* null mutants), inhibition of ventral muscles induced by iMN activation was significantly attenuated compared to wildtype animals (Figures 4D and 4E, Wildtype and GABA_A_-). In contrast, iMN activation-mediated inhibition on the dorsal muscles was only modestly reduced (Figures 4D and 4E, Wildtype and GABA_A_-). Thus the GABA_A_ receptor is critical for inhibition to ventral muscles, and less so for dorsal muscles.

In contrast, in the absence of GABA_B_ receptors (*gbb-2* null mutants), iMN activation inhibited ventral muscle to the same degree as in wildtype animals, while inhibition of dorsal muscle was significantly attenuated (Figures 4D and 4E). The predominant requirement of an extrasynaptic GABA_B_ receptor for dorsal muscle inhibition, together with the absence of NMJs from iMNs onto dorsal muscles (Figures 1B, left panel, and S1), indicate a predominant role of extrasynaptic GABA signaling in mediating dorsal muscle relaxation.

### Extrasynaptic GABA inhibition may promote dorsal muscle relaxation through multiple cells

To further examine through which cells extrasynaptic GABA inhibition contributes to dorsal muscle relaxation, we examined where GABA_B_ receptors reside in L1 larvae. GABA_B_ receptors function as an obligatory heterodimer for membrane trafficking across animals ([40–42] and in adult *C. elegans* [35–37]). We examined the expression patterns of these subunits, GBB-1 and GBB-2 using fosmid reporters (Method).

GABA_A_ receptors inhibit ventral muscle synaptically, consistent with their tightly clustered expression restricted to ventral muscles (Figure 4C, top panel). By contrast, the dorsal muscle-inhibiting GABA_B_ subunits are not restricted to dorsal muscles. GBB-1::GFP signal is weak, with the most prominent presence in the motor neuron soma along the ventral cord (Figure 4C, middle panel). GBB-2::GFP signal is stronger, detectable in the motor neuron soma and neurite along the ventral cord, as well as in dorsal and ventral muscles (Figure 4C, bottom panel). Weak GBB-1::GFP and stronger GBB-2::GFP signals are also present in some neuron soma in the head (Figure 4C, middle and bottom panels).

Such expression patterns suggest that multiple cells may contribute to extrasynaptic GABA inhibition-mediated dorsal muscle relaxation, from inhibition of dorsal muscles themselves, to inhibition of dorsal muscle-activating eMNs, even the unidentified upper layer interneurons. Among these cells, motor neuron somas exhibit the most prominent GABA_B_ reporter overlap, and also reside in the closest proximity to iMN’s GABA releasing NMJs. Indeed, optogenetic activation of iMN motor neurons led to strong inhibition of the dorsal muscle-innervating eMN motor neurons (Figure 4B, Wildtype), and the inhibition was attenuated, though not abolished in *gbb-2* null mutants (Figure 4B, GABA_B_-). Thus, extrasynaptic GABA signaling likely promotes dorsal muscle relaxation through multiple mechanisms, from direct inhibition of dorsal muscles, to indirect inhibition through neurons that drive dorsal muscle contractions. The latter includes eMN motor neurons, and we do not exclude potential involvement of other neurons. A broader cellular source of inhibition is consistent with the less constrained nature of extrasynaptic signaling.

Inhibition of dorsal muscles was not abolished in *gbb-2* null mutant larvae. Only in L1 larvae that lack both GABA_A_ and GABA_B_ receptors, iMN activation failed to fully inhibit dorsal as well as ventral muscles (Figures 4D and 4E, GABA_A_-;GABA_B_-). Mutant L1 larvae that do not synthesize GABA (*unc-25* null mutants) recapitulated this response (Figure S4). Importantly, despite making a minor contribution to the extent of dorsal muscle inactivation, the GABA_A_ receptors controlled the rate of inhibition in not only ventral, but also dorsal muscles (Figure 4F, GABA_A_-). Excitatory motor neurons are proprioceptive in the adult motor circuit^14^. Thus GABA receptor-mediated relaxation of ventral muscles and subsequent reduction of body bending may further contribute to the inhibition of dorsal muscle-innervating eMNs. Consistent with this notion, iMN-mediated inhibition of excitatory motor neuron was reduced but not abolished in the absence of GABA_B_ (Figure 4B).

Together, these results demonstrate that the iMNs indirectly relax dorsal muscles, likely through multiple mechanisms. Due to the eMN’s proximity to released GABA, expression of both GBB-1 and GBB-2 subunits, and proprioceptive gating, we propose that an extrasynaptic inhibition to the eMNs work in concert with the synaptic and extrasynaptic inhibition of ventral muscles to promote an exit after dorsal bending.

Therefore, in L1 larvae, motor neurons form a circuit where synaptic wiring drives dorsal bending. This circuit regulates its own exit from dorsal bending through negative feedback, prompted by extrasynaptic GABA signaling.

### Ventral muscle excitation requires cholinergic premotor interneurons

A motor circuit to drive dorsal bending and its own exit does not produce ventral bending. Ventral bending requires ventral muscle contraction that occurs in synchrony with dorsal muscle relaxation. Our complete EM reconstruction of juvenile L1 larvae only revealed NMJ inputs from iMN motor neurons (Figure S1; [25]), which we demonstrated to be inhibitory (above). Other mechanisms must excite ventral muscles in order for the L1 motor circuit to transit from dorsal to ventral bending.

To reveal this mechanism, we first considered the simplest possibility: a higher myogenic activity of ventral muscles compensates for the absence of motor neuron excitation. However, we found that optogenetic inhibition of the L1 larva’s entire nervous system led to similarly silenced dorsal and ventral muscles (Figure S5A). This shows that ventral muscle activity is not myogenic, but is driven by neurons.

To identify neurons that excite ventral muscles, we turned to optogenetic stimulation. Because ventral muscles receive inhibitory NMJs from iMNs, in wildtype larvae, pan-neuronal excitation activated dorsal muscles but inhibited ventral muscles (Figure S5C, left panel). In L1 larvae that do not synthesize GABA (*GAD/unc-25* null mutants), however, we observed activation of both ventral and dorsal muscles (Figure S5C, right panel).

We then applied similar optogenetic manipulation to sub-groups of neurons in wildtype and *unc-25* larvae to identify the neuronal source for ventral muscle activation. We found that optogenetic manipulation of all cholinergic neurons recapitulated the effects of pan-neuronal manipulation: pan-cholinergic neuron inhibition abolished activity in both dorsal and ventral muscles (Figure S5B). Pan-cholinergic neuron stimulation led to dorsal muscle excitation and ventral muscle inhibition in wildtype larvae (Figure S5D, left panel), but activation of both dorsal and ventral muscles in *unc-25* mutant larvae (Figure S5D, right panel). Increased ventral muscle activity upon removal of GABA is consistent with our previous result that cholinergic motor neurons activate iMNs to relax ventral muscles. Thus, cholinergic neurons are responsible for ventral muscle activation.

We have shown by EM reconstruction that cholinergic eMNs do not make NMJs to ventral muscles (Figure S1) and they activate dorsal muscles (Figure 2). However, premotor interneurons (eINs) are cholinergic^43^, their axons span the ventral nerve cord, making them the candidate source of ventral muscle excitation. Along the nerve cord, two eINs, AVA and PVC, make numerous synapses to all eMNs. Others (AVB, AVE, AVD) make synapses to AVA and PVC, and to one another (Figure 1B, left panel).

We found that optogenetic inhibition of eINs by GtACR2 silenced both dorsal and ventral muscles (Figure 5A; Video S5A). In mutant larvae without GABA (*unc-25*), optogenetic stimulation of eINs activated both dorsal and ventral muscles, with a stronger activation of ventral muscles (Figure 5B; Video S5B). Patterns of dorsal and ventral muscle activity, as in *unc-25* mutants (Figure 3F), differed: dorsal activity was in correlation with local curvature, whereas ventral activity was uniform along the body (Video S5B). These results suggest that ventral muscles may be uniformly activated by extrasynaptic cholinergic signaling from the eINs.

**Figure 5.**
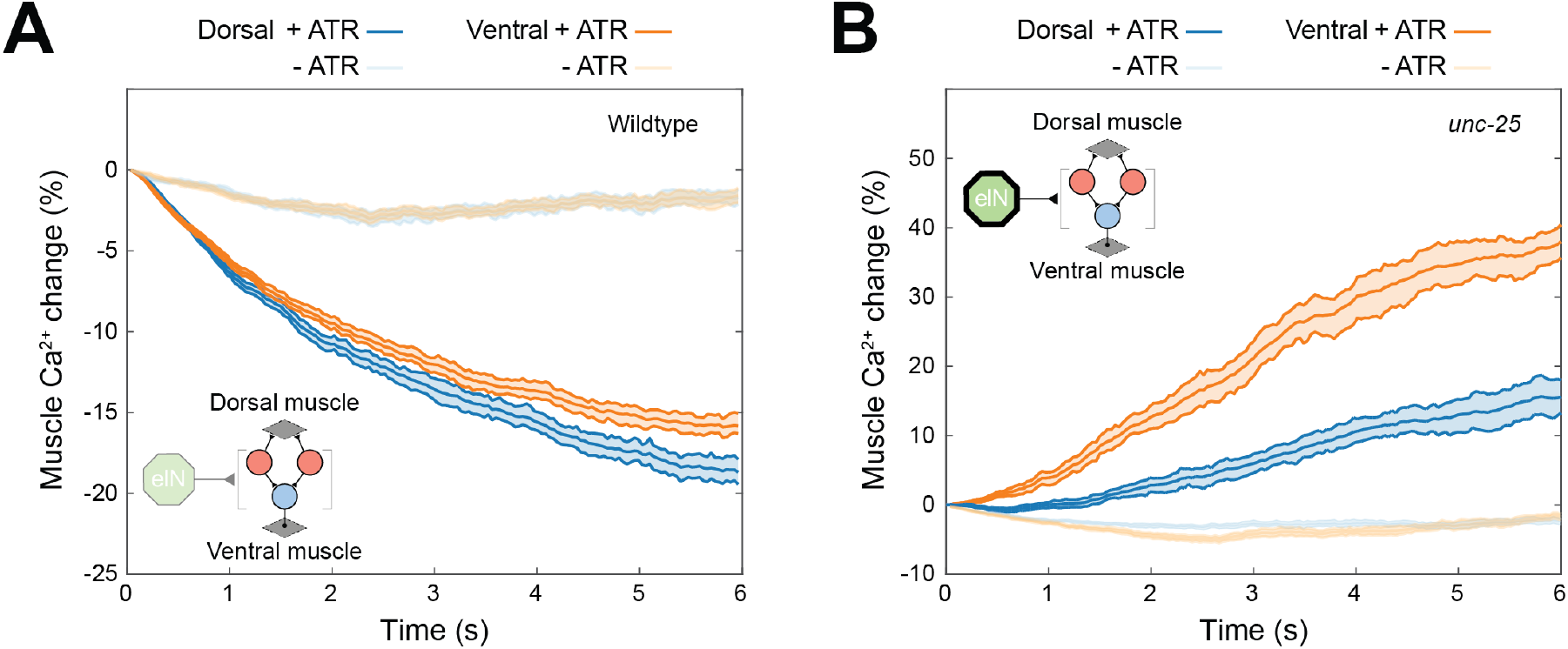
Cholinergic eINs underlie ventral muscle contraction. (A) (Inset) Schematic of simultaneous inactivation of all eINs and muscle calcium imaging in wildtype L1 larvae. Y-axis plots percentage changes of the muscle activity from *t* = 0. Inhibition of eINs by GtACR2 reduced both dorsal and ventral muscle activity in wildtype larvae. Control group(-ATR): 28 stimulation epochs from 6 larvae; Experimental group(+ ATR): 49 stimulation epochs from 13 larvae. (B) (Inset) Schematic of simultaneous eIN optogenetic activation and muscle calcium imaging in *unc-25* L1 larvae. Y-axis plots percentage changes of the muscle activity from *t* = 0. Activation of eINs by Chrimson elevated both dorsal and ventral muscle activity in *unc-25* larvae. Control group(-ATR), 51 stimulation epochs from 17 larvae; Experimental group(+ ATR): 43 stimulation epochs from 12 larvae.

To further determine whether eINs are required for ventral muscle activation, we ablated them using miniSOG. The eIN-ablated wildtype L1 larvae exhibited diminished ventral muscle activity, but with residual dorsal muscle activity (Figure 6A; Videos S6A and S6E). When the inhibitory input to ventral muscles was removed in the *unc-25* mutant larvae, ablation of eINs also reduced ventral muscle, but not dorsal muscle activity (Figure 6B; Videos S6B and S6F). Reduced ventral muscle activity in both wildtype and *unc-25* mutant larvae confirms that eINs are required for ventral muscle excitation. Persistent dorsal muscle activity is consistent with the presence of endogenous eMN activity, as previously observed in the adult circuit^13,16^. These results establish the requirement of eINs in stimulating ventral muscles.

**Figure 6.**
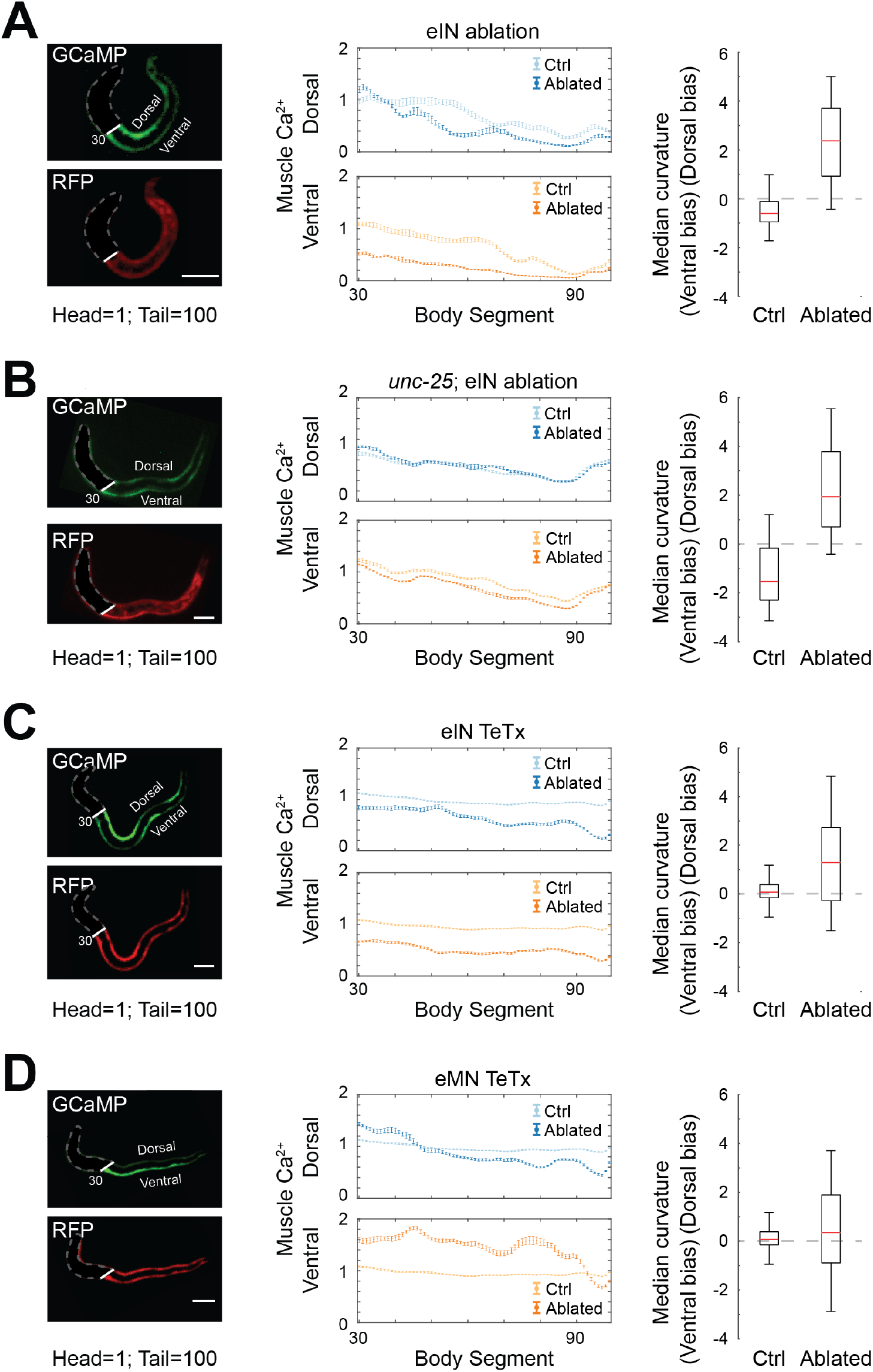
Extrasynaptic transmission from eINs is required for ventral bending. (A) Muscle activity and body curvature of L1 larvae without eINs. (Left panels) Example images of muscle GCaMP::RFP. (Middle panels) Average dorsal (top) and ventral (bottom) muscle activity in the control (lighter colors) and experimental group (darker colors). The control and eIN-ablated groups were the same set of animals expressing miniSOG in eINs, either without or with exposure to blue light prior to calcium imaging (Methods). (Right panel) Median curvature across the body (segments 33-95) of eIN ablated group (Ablated) compared with the control group (Ctrl). eIN ablation led to stronger reduction of ventral activity; these larvae exhibited a bias for dorsal bending along the body. N = 10 (Ctrl) and N = 11 (Ablated) larvae. (B) Same as (A), but for mutant L1 larvae without GABA *GAD/unc-25(e156)*. Upon eIN ablation, L1 larvae exhibited reduced ventral muscle activity and a bias for dorsal bending across the body. N = 13 (Ctrl) and N = 15 (Ablated) larvae. (C) Muscle activity and body curvature of L1 larvae where synaptic vesicle release from eINs was constitutively blocked by TeTx. (Left panels) Example images of muscle GCaMP::RFP. (Middle panels) Average dorsal (top) and ventral (bottom) muscle activity of the L1 larvae expressing TeTx in eINs (darker colors). The control group (Ctrl) were L1 larvae not expressing TeTx (lighter colors). (Right panel) Median curvature across the body (segments 33-95) of synaptic vesicle release blocked group (TeTx) compared with the control (Ctrl) group. Blocking vesicle release from eINs also resulted in a reduction in ventral muscle activity and a bias for dorsal bending across the body. N = 10 (Ctrl) and N =14 (TeTx) larvae. (D) Same as (C), but with L1 larvae expressing TeTx in eMNs. The control group were L1 larvae not expressing TeTx (Ctrl; lighter colored lines in the middle panels). Blocking synaptic vesicle release from eMNs resulted in a higher ventral muscle activity (middle panels), but not a consistent bias towards either ventral or dorsal bending across the body (right panel). N =10 (Ctrl) and N = 12 (TeTx) larvae.

### Cholinergic premotor interneurons activate ventral muscles via extrasynaptic signaling

Our EM reconstruction of L1 larvae have shown that only iMNs make synapses to ventral muscles (Figure 1B, left panel, and S1). Because eINs do not make synapses to ventral muscles (Figures 1B, left panel, and S1), they may activate ventral muscles either by ephaptic coupling, or by extrasynaptic mechanisms that require neurotransmitter release.

We distinguished these two possibilities by assessing the effect of blocking synaptic vesicle fusion using tetanus toxin (TeTx)^44^. When vesicular release from all eINs was blocked, activity in both dorsal and ventral muscles was reduced (Figure 6C; Videos S6C and S6G). Vesicular release is needed for eINs to activate dorsal muscles because they innervate eMNs, as expected. Reduced ventral muscle activation, however, demonstrates that neurotransmitter release is also needed for eINs to activate ventral muscles. Similar to effects of eIN ablation, ventral muscle activity was more severely reduced than dorsal muscles (Figure 6C, middle panels). Lastly, RNAi-mediated knockdown of *cha-1*—a gene required for acetylcholine synthesis^45^—from all eINs also led to more severely reduce activity in ventral muscles (Figure S6A). These results argue for the requirement of vesicular release of acetylcholine to drive ventral muscle activation. Extrasynaptic accumulation of acetylcholine is also consistent with the uniform increase of ventral muscle activity during eIN stimulation.

Together, these results suggest that cholinergic eINs stimulate ventral muscles through extrasynaptic acetylcholine accumulation.

### L1 motor circuit orchestrates dorsal-ventral symmetry by anti-phasic entrainment

Our data demonstrated that synaptic wiring among motor neurons is built to make only dorsal bends. But with ventral muscles tonically activated by eINs, the same wiring can entrain ventral muscles to generate complementary ventral bends. An implication of this model is that ventral muscle excitation takes places independently of motor neurons, but synaptic wiring among motor neurons is required to generate coordinated dorsal-ventral bending.

Consistent with this implication, blocking synaptic transmission from eMNs and eINs by TeTx led to opposite effects on dorsal and ventral muscles (Figure 6D). Eliminating eINs or their synaptic output resulted in dorsal muscle activity being higher than ventral muscle activity (Figures 6A, 6B, and 6C; Videos S6A, S6B, and S6C). Eliminating eMN’s synaptic output led to ventral muscle activity being higher than dorsal muscle activity (Figure 6D; Video S6D).

Behaviorally, however, only dorsal muscle’s calcium signals predict a crawling larva’s bending pattern, reflecting direct input from eMNs. When eINs or their synaptic output were eliminated, regardless of the absolute level of residual dorsal activity, they led to a persistent bias for dorsal bending across body segments, in wildtype as well as *unc-25* mutant larvae (Figures 6A, 6B, and 6C, right panels; Videos S6E, S6F, and S6G).

When eMN synaptic output was blocked, regardless of a high ventral muscle activity (Figure 6D, left and middle panels), L1 larvae exhibited no consistent dorsal or ventral bending bias across body segments (Figure 6D, right panel; Video S6H). This coincides with a spatially uniform calcium signal in their ventral muscles that lacked correlation with curvature (Figure 6D, left panel; Video S6D). The lack of correlation is similar to the calcium pattern of ventral muscles in the *unc-25* mutant larvae (Figure 3F).

### A computational model: generating a symmetric motor pattern with an asymmetric circuit

How bending is generated in each segment is illustrated in Figure 7A. In L1 larvae, dorsal and ventral muscles have similar excitability (Figure S5). Rhythmic dorsal bending is directly mediated by oscillatory activity of cholinergic motor neurons. Cholinergic motor neurons simultaneously activate GABAergic neurons that relax ventral muscles. The net effect is a dorsal bend (Figure 7A, i).

**Figure 7.**
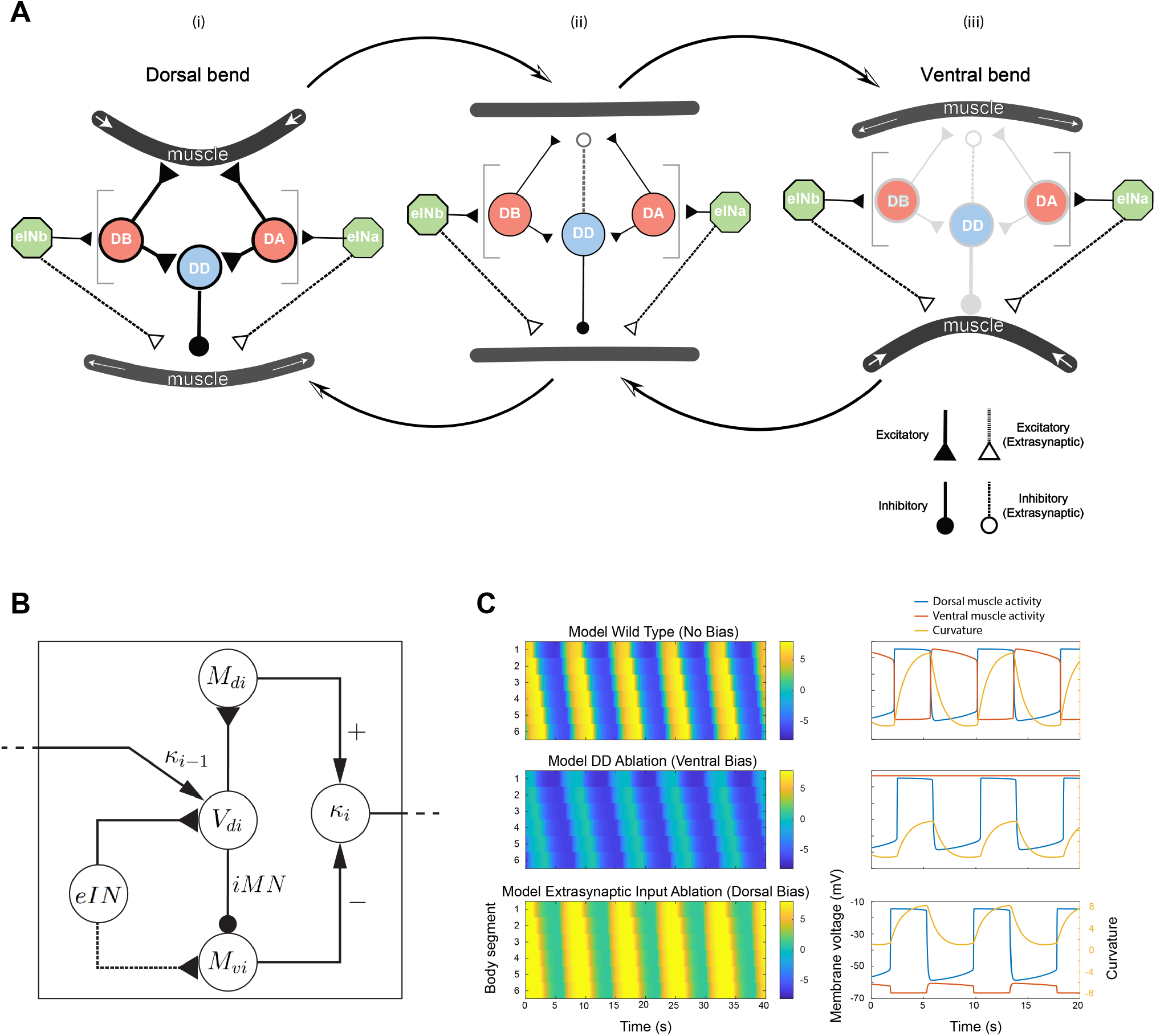
A computational model of the L1 motor circuit. (A) L1 larvae integrate synaptic transmission and extrasynaptic transmission to generate alternating body bends. Schematic depicts transitions between three bending phases. (i) Synaptic wiring from the eMN and iMN contracts dorsal muscle and relaxes ventral muscle, generating a dorsal bend. (ii) Reduced synaptic transmission from the eMN and inhibitory feedback from the iMN reduces dorsal contraction and ventral relaxation, promoting exit from a dorsal bend. (iii) Further reduction of eMN activity removes inhibition on ventral muscle, allowing extrasynaptic excitation from the eIN to generate a ventral bend. (B) Schematic of the computational model at one body segment. *V*_*di*_, *M*_*di*_, *M*_*vi*_ denote membrane potentials of the eMN, dorsal muscle and ventral muscle at segment *i*, respectively. *κ*_*i*_ is the curvature at segment *i. κ*_*i*−1_ represents the proprioceptive input from the anterior segment. *DD* denotes the implicit inhibitory synapse on ventral muscle. Inverted and circular arrowheads denote excitatory and inhibitory synapses, respectively. Dotted arrow denotes extrasynaptic excitation. Regular arrowheads represent mathematical operations. (C) (Left) Curvature kymographs from simulations in different conditions. (Right) Time-series of dorsal-ventral muscle activity and curvature at mid-body (segment 3). (Top) Simulation of the full circuit produces traveling waves along the body. (Middle) Simulation of the circuit without inhibitory synapse produces traveling waves only in dorsal muscles. (Bottom) Simulation of the circuit without extrasynaptic input from the eIN leads to severe reduction of ventral muscle activity and strong bias for dorsal bending.

Rhythmic ventral bending is not mediated by its own oscillator, but by anti-phasic entrainment to the oscillator that drives dorsal bends. Ventral muscles are uniformly excited by extrasynaptic acetylcholine from premotor interneurons. This permits ventral bends to occur when and where the dorsal oscillator’s activity is low (Figure 7A, iii).

Transitions between dorsal bending and ventral bending are also facilitated by negative feedback within each segment: GABA released by inhibitory motor neurons relaxes ventral muscles and extrasynaptically inhibits excitatory motor neurons that drive dorsal bending (Figure 7A, ii). Thus, in L1 larvae, the same set of motor neurons drive both dorsal and ventral bends. This is in contrast to the adult, which has two symmetrically wired motor subcircuits that separately drive dorsal and ventral bends.

We developed a phenomenological model that describes the L1 motor circuit dynamics (Methods; Figure 7B). Similar to previous work in the adult motor circuit^16^, the eMN is modeled as an oscillator (*V*_*di*_) with calcium and potassium conductances. It provides direct input to dorsal muscles (*M*_*di*_) through an excitatory synapse. The eMN relays a copy of this oscillation to ventral muscles (*M*_*vi*_), inverting its sign through an inhibitory synapse (*DD*). When dorsal and ventral muscles have the same resting membrane potential, the inhibitory synapse rectifies dorsal oscillations. This produces alternating dorsal-ventral bends with a deeper dorsal bend (Figure 7C, lower panel). An extrasynaptic excitatory input to ventral muscles (*eIN*) alleviates the dorsal bias (Figure 7C, upper panel).

We modeled propagating bending waves through proprioceptive input between adjacent eMNs (*κ*_*i*_). As in previous models^16,46^, directional wave propagation is encoded by proprioceptive input where the eMN in segment *i* receives curvature-dependent input from *i* − 1 (for forward locomotion) or from *i* + 1 (for backward locomotion). Net bending is calculated from the extent of dorsal and ventral muscle activation in each segment (Methods). We obtained parameters by fitting the model outputs to key experimental findings on dorsal and ventral output symmetry (Figure 6).

Without the inhibitory synapse to ventral muscles, this model predicts stalled wave propagation on the ventral side. This fully recapitulates the lack of correlation of ventral muscle activity with bending of GABA- (*unc-25*) mutant larvae (Figure 3F). Perturbation of the inhibitory synapse in this model generates bending with a ventral bias across the body. Behaviorally, while *unc-25* mutants exhibit a ventral bias, they do also generate dorsal bends (Videos S7A and 7B). Our model can recapitulate this phenomenon by adjusting the extrasynaptic input to ventral muscles. This suggests that premotor interneurons may receive a feedback from the downstream motor circuit.

Elimination of extrasynaptic input in this model generates dorsal bending across the body, which recapitulates the behavior of eIN-ablated or blocked larvae (Figures 6A, 6B, and 6C). Compared to a simpler model, where ventral muscles are spontaneously active, an extrasynaptic input to ventral muscles confers more flexibility for bending waves. This functional configuration offers a temporary solution for symmetric bending before the motor circuit matures.

## Discussion

Symmetric motor circuit inputs are assumed to be required for symmetric motor outputs^3–5^. *C. elegans* is born with only motor neurons that are wired analogously to one motor subcircuit for dorsal bending in adults. Over the course of larval development, extensive neurogenesis and rewiring adds another motor subcircuit for ventral bending and eliminates asymmetries in the layout of the newborn circuit.

The larva and adult generate the same motor pattern despite these profound anatomical differences. *C. elegans* reveals a strong drive to maintain a stable and symmetric motor output throughout development. We show that L1 larvae achieve its alternating dorsal-ventral motor pattern by using its adult-like motor circuit for dorsal bending to also anti-phasically entrain its ventral bending.

### Summary of results

By EM reconstruction of juvenile L1 larvae prior to post-embryonic neurogenesis, we show that dorsal muscles receive NMJs from cholinergic motor neurons whereas ventral muscles receive NMJs from GABAergic motor neurons. There are no significant morphological synapses, chemical or electrical, from other cells to these muscles. By functional imaging and optical stimulation, we demonstrate that the NMJ inputs to dorsal muscles are excitatory and the NMJ inputs to ventral muscles are inhibitory, thus forming a circuit for dorsal bending.

The absence of wired inhibitory synapses to dorsal muscles and excitatory synapses to ventral muscles raises the possibility of extrasynaptic signaling or ephaptic coupling to compensate for wired inputs on both sides of the animal. By analyzing genetic mutations that separately disrupt synaptic and extrasynaptic GABA signaling, we found that extrasynaptic GABA signaling provides the inhibitory inputs to dorsal muscles. By optogenetic activation and silencing of muscles and different groups of neurons, we found that ventral muscle activation requires the excitatory premotor interneurons, independent of their wired inputs to the excitatory dorsal-muscle innervating motor neurons. By anatomically ablating these premotor interneurons, blocking their vesicular release, or reducing their vesicular load of acetylcholine, we showed that extrasynaptic cholinergic signaling from these premotor interneurons uniformly activates ventral muscles. With tonically potentiated ventral muscles, the wired motor neuron circuit that drive dorsal bending can produce spatially coordinated, anti-phasic ventral bending.

With these results, we built and tested a model where the synaptic transmission and extrasynaptic transmission work in concert to enable the larva to produce a adult-like, undulatory motor pattern before anatomical maturation of the motor circuit. This model validates the adequacy of the larval solution for generating motor patterns, and reveals the limitation of this strategy.

### Circuit degeneracy maintains an animal’s motor patterns

The *C. elegans* motor circuit is a profound example of circuit degeneracy where similar functional outputs can have multiple network solutions^21–23^. Other examples include the similar motor patterns exhibited by two molluscs with substantially different functional connectivity^7^, and maintenance of a pyloric circuit’s output at different temperatures by varying parameters of synaptic connections^47^. Circuit degeneracy has been proposed to play a critical role in population diversity, adaptability to changing environments, and species evolution.

Our results demonstrate that degeneracy also minimizes functional disruption to locomotion during development, representing an intrinsic drive to maintain stable motor output in response to ecological demand.

### Extrasynaptic signaling enables an asymmetric L1 motor circuit to generate symmetric motor output

L1 larva’s alternative solution for dorsal-ventral bending requires extrasynaptic signaling at multiple layers of its motor circuit.

Dorsal bending is driven by a configuration that resembles the original half-center model proposed for flexor-extensor coordination during limb movement^1^: synaptic wiring of excitatory and inhibitory motor neurons coordinates dorsal muscle excitation with ventral muscle relaxation.

In contrast, ventral bending is generated by anti-phasic entrainment to the rhythmic activity of this dorsal half-center. For this to occur, it is necessary for ventral muscles to be uniformly activated. Acetylcholine from premotor interneurons, which accumulates and acts extrasynaptically, fulfills this role.

Intrinsic oscillatory activities of excitatory motor neurons underlie transitions from dorsal to ventral bending, while GABA release from inhibitory motor neurons facilitates and modulates this transition. Acting extrasynaptically, this negative feedback inhibits excitatory motor neurons and therefore promotes dorsal muscles to relax after contraction.

Thus extrasynaptic signaling is necessary for the asymmetric motor circuit to create symmetric output.

### Extrasynaptic signaling as an adaptive strategy for a developing motor circuit to create mature behavior

Extrasynaptic accumulation of fast neurotransmitters may result from vesicular release at extrasynaptic sites, or by extracellular diffusion of synaptically released neurotransmitters^48^. We favor the diffusion mechanism for extrasynaptic activation of ventral muscles, because we did not observe an overt non-synaptic vesicle accumulation in the EM volume of newborn L1 larvae.

Extrasynaptic signaling is widely observed in other systems^49–52^. It is one mechanism for neuromodulation, which allows hard-wired mature circuits to modify and adapt their output to different contexts^53,54^. Our results reveal that during *C. elegans* early development, extrasynaptic transmission is essential, functionally compensating for the absence of an entire motor subcircuit. As larvae mature, this role is taken over by postembryonic motor neurons and synaptic rewiring^25^.

Not all extrasynaptic transmission is replaced as the motor circuit matures. Extrasynaptic feedback from GABAergic motor neurons to excitatory motor neurons persists in the adult^35–37^. It may play similar roles in both dorsal and ventral motor subcircuits.

Extrasynaptic mechanisms may work particularly well in the larva because of its small size. In a short-lived animal, acquiring full mobility at birth is of probable fitness benefit. Extrasynaptic signaling thus represents an adaptive strategy for development.

### Neurons adopt new functions to compensate for structural changes

In small circuits, individual neurons often adopt multiple functions to expand their behavioral repertoires^55,56^. Multi-functionality is generally considered to be a property of mature neural circuits.

In the adult, cholinergic motor neurons are rhythm generators for forward and backward movement^13,15,16^. Cholinergic premotor interneurons signal to switch the these two modes of locomotion^16,19^. Our calcium imaging of premotor interneurons (Figure S6B) and behavioral effects upon their activation (Figure S6C) showed that this role is likely conserved across development. However, premotor interneurons assume an additional role in the L1 larva: they directly activate ventral muscles. This excitation is independent of motor neurons. Neither is this excitation the consequence of positive feedback from cholinergic motor neurons, because cholinergic motor neurons specifically activated dorsal muscles.

The essential role of premotor interneurons in ventral muscle excitation is likely temporary: the expression pattern of acetylcholinesterase encoding genes, whose enzymatic activity determines extrasynaptic acetylcholine buildup, exhibits dorsal-ventral asymmetry in the newborn larva. Asymmetry disappears after the L1 stage, coinciding with the emergence of ventral muscle-innervating motor neurons^57^.

Thus, a neuron’s flexibility to adopt multiple roles also provides a means to compensate for a circuit’s developmental immaturity.

### An adequate but temporary solution for a developing motor circuit

How L1 larvae generate a symmetric motor pattern with an immature, asymmetric circuit has been an enigma. Several hypotheses have been proposed^58,59^, but our experimental findings are inconsistent with these models.

Prior models assumed ventral bending to be myogenic. Our model incorporates a neural source of ventral muscle excitation established by experimental findings, providing a more flexible solution. Because the relative strength of synaptic and extrasynaptic inputs from premotor interneurons — the source of ventral muscle excitation — can be adjusted separately, this model confers flexibility and a range of possible bending patterns with balanced or imbalanced dorsal-ventral bends.

In this L1 model, dorsal and ventral muscles are driven by the same oscillators. Consequently, their activation is not fully independent of each other. This implies that synaptic input to dorsal muscles alone determines wave propagation. This prediction recapitulates the experimental outcome of decoupling of dorsal and ventral muscle activity: when we removed synaptic output of dorsal oscillators, or the inhibitory synapse that relays this oscillation to ventral muscles, ventral calcium signals did not propagate and did not correlate with bending.

This configuration differs from the adult, where both dorsal and ventral muscles are driven by dedicated motor subcircuits. Differential activation of dorsal and ventral muscles might have advantages in the worm’s natural habitat, where it would enable more robust exploration in a 3D environment^60^. Thus, L1 larva’s configuration offers an adequate but temporary solution.

## Star Methods

### Key resources table

| REAGENT or RESOURCE | SOURCE | IDENTIFIER |
| --- | --- | --- |
| **Antibodies** |  |  |
| RFP antibody [5F8] | Chromotek | Cat#5f8-20 |
| GFP monoclonal antibody | Invitrogen | Cat#A-11120 |
| **Bacterial strains** |  |  |
| *E. coli* | *Caenorhabditis* Genetics Center | RRID: WB-STRAIN:OP50-1 |
| **Experimental models** |  |  |
| *C. elegans* | This paper | Table S2 |
| Recombinant DNA | This paper | Table S2 |
| **Software and algorithms** |  |  |
| ImageJ | NIH Image | RRID: SCR_003070 |
| NIS elements | Nikon | RRID: SCR_014329 |
| MATLAB | MathWorks | RRID: SCR_001622 |
| GraphPad Prism | GraphPad | RRID: SCR_002798 |
| **Codes** |  |  |
| Github | This paper | https://github.com/zhen-lab/Beta_Function_Analysis |
| **Reader and Reagent Guides** | This paper | Table S1, Table S2 |

## RESOURCE AVAILABILITY

### Lead Contact

Further information and requests should be directed to lead contacts.

### Materials Availability

Constructs and strains generated in this study will be deposited at Addgene and CGC for distribution, and also available upon request.

### Data and Code Availability

All MATLAB scripts developed for behavioral analyses, neuronal and muscle calcium imaging analyses, and modeling are available in GitHub https://github.com/zhen-lab/Beta_Function_Analysis.

## EXPERIMENTAL MODEL AND SUBJECT DETAILS

*C. elegans* strains were grown and maintained on nematode growth media (NGM) plates seeded with the *Escherichia coli* strain OP50 at 20-22.5^°^C.

## METHOD DETAILS

### Construction of transgenic strains

Transgenic ZM strains were generated by injecting fosmid or plasmid constructs (pJH) at 2-50ng/ul, with respective injection markers into indicated genetic backgrounds, to produce those with extra chromosomal arrays (hpEx). They were integrated into the genome by UV irradiation of the Ex lines followed by selective screening and outcrossing against N2 wild-type strains (hpIs). Other transgenic arrays or strains were acquired from either *CGC* or individual laboratories. A reader and reagent guide, which provides detailed information and usage of constructs and strains are listed in two tables below.

**Table S1:** Nomenclature of cells, promoters, and promoter expression patterns

| Neuron group abbreviations | Neurons included | Promoters used | Overlap in neuron targets by promoters in L1 |
| --- | --- | --- | --- |
| Cholinergic premotor interneurons (eIN) | AVA, AVB, AVE, AVD, PVC | *Pnmr-1*<br>*Psra-11*<br>*Ptwk-40s*<br>*Prig-3LoxP/Ptwk-40Cre*<br>*Prig-3LoxP/Psra-11/twk-40Cre*<br>*Plgc-55s* | AVA, AVE, AVD, PVC (+ others)<br>AVB<br>AVA, AVE, AVB<br>AVA<br>AVA, AVB<br>AVB |
| Excitatory motor neurons (eMN) | DA, DB | *Pacr-2s*<br>*Pacr-5*<br>*Punc-4* | DB, DA (weak)<br>DB<br>DA |
| Inhibitory motor neurons (iMN) | DD | *Punc-25*<br>*Pttr-39* | DD<br>DD |
| Panneuronal | All neurons | *Prgef-1* | All neurons |
| Pan-cholinergic | All cholinergic neurons | *Punc-17* | All cholinergic neurons |

**Table S2.**
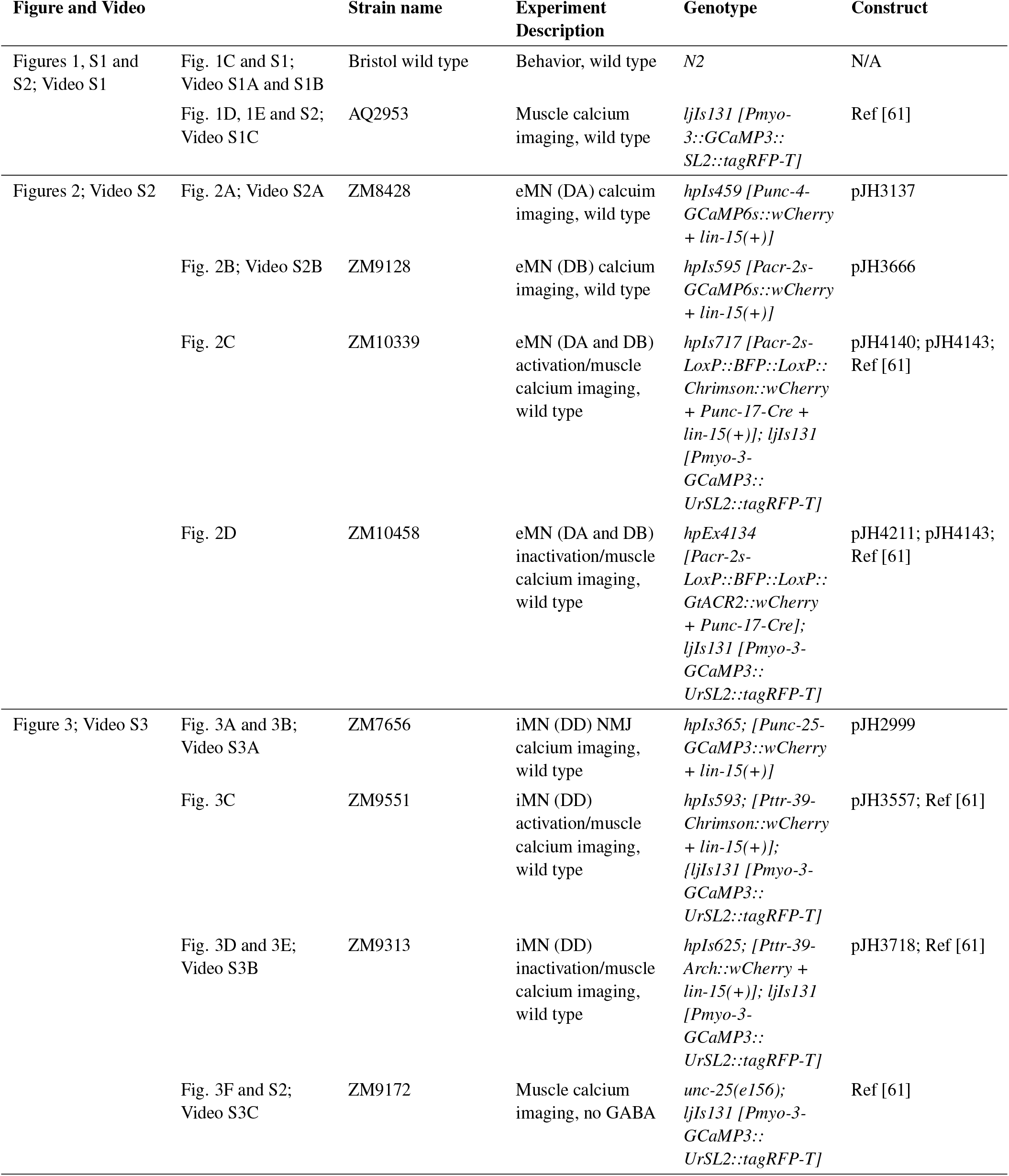
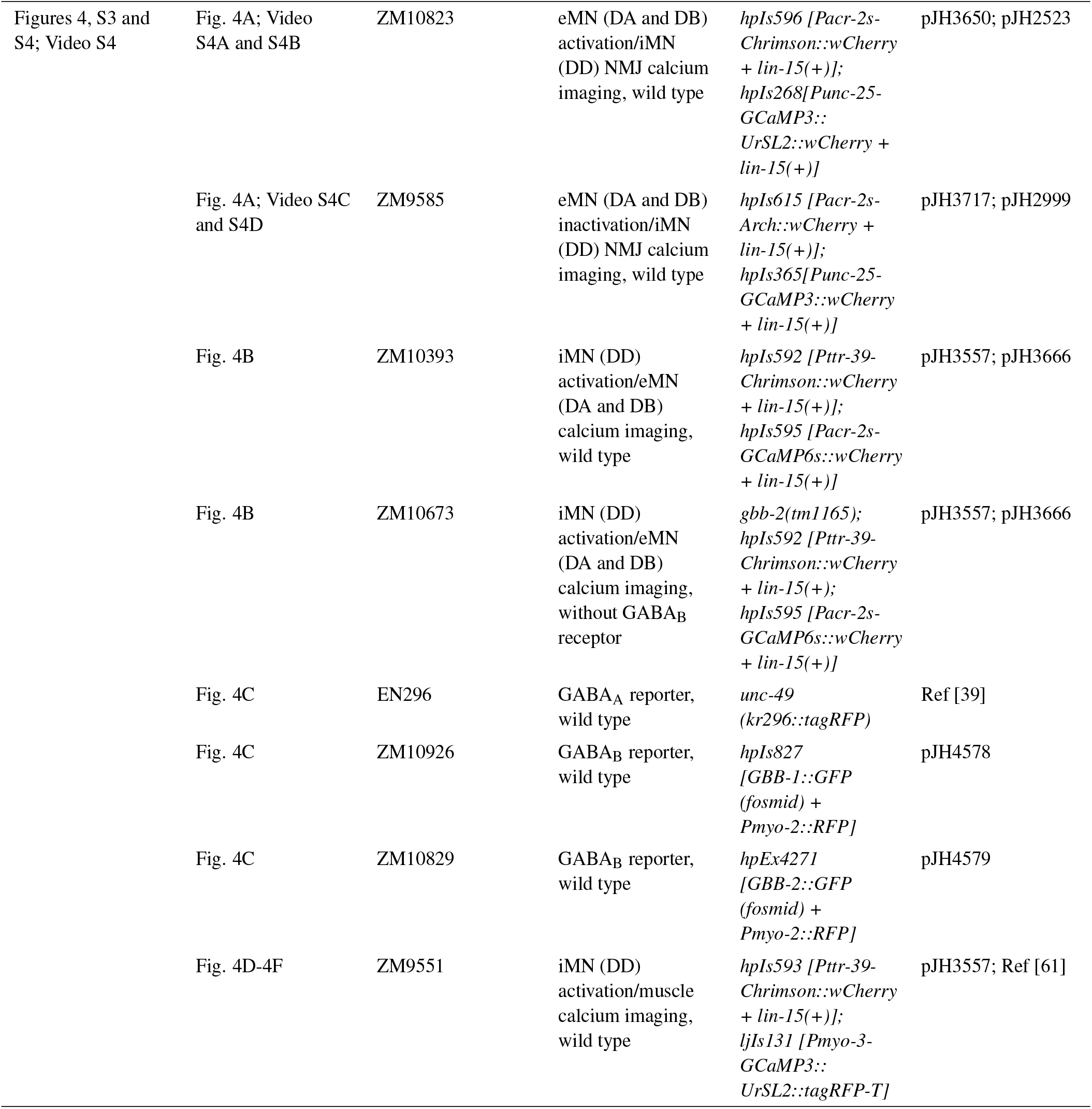
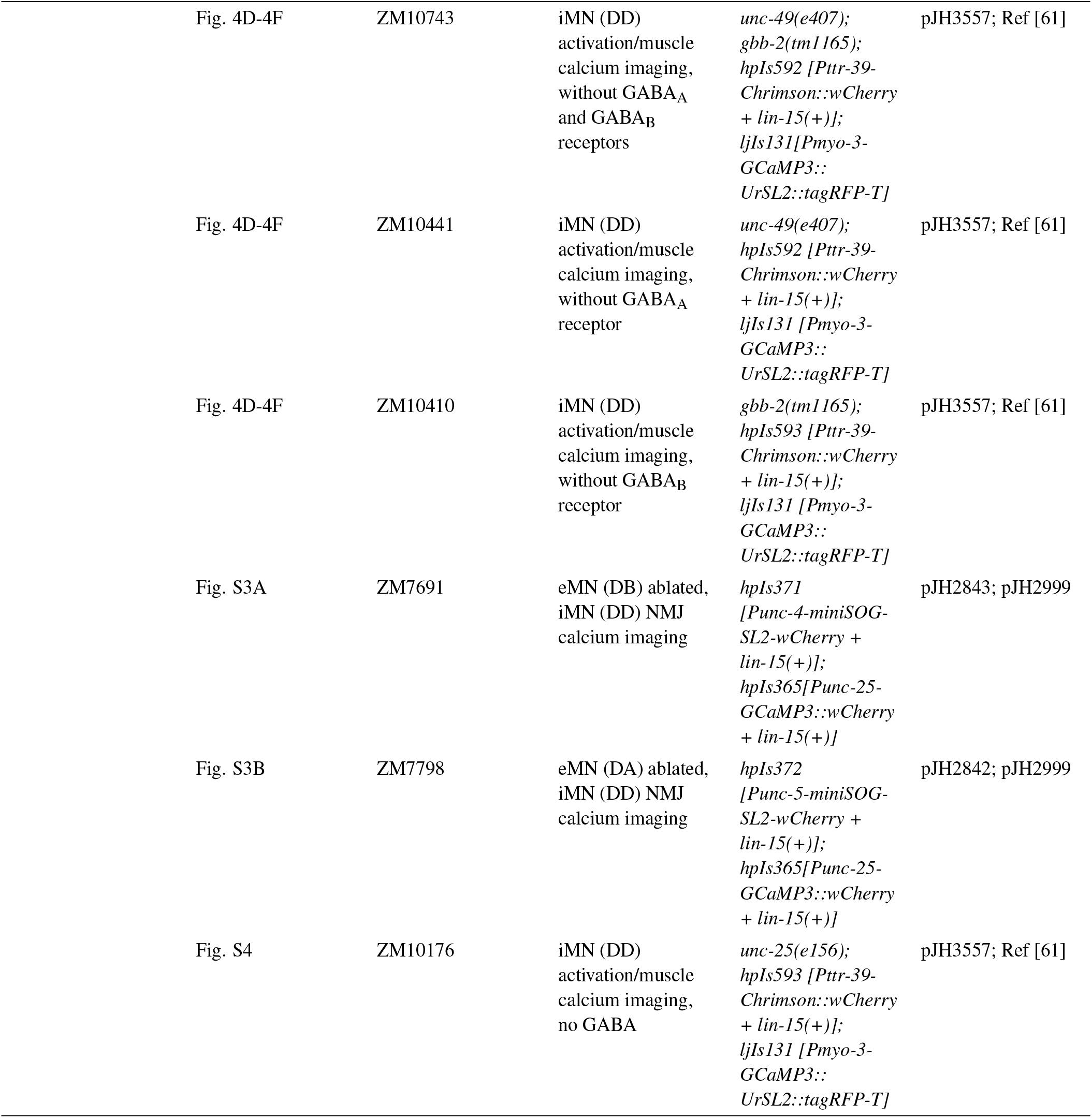
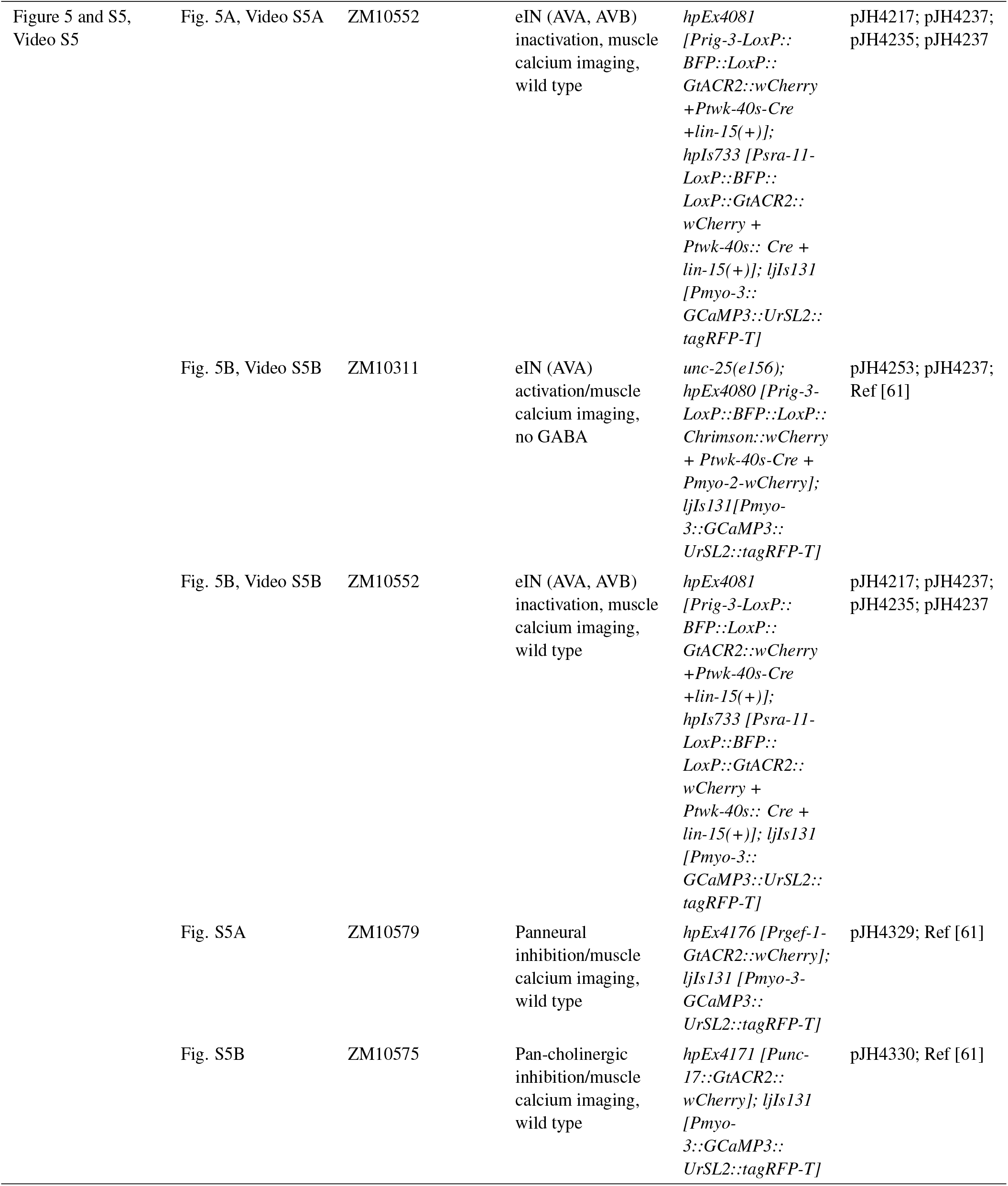
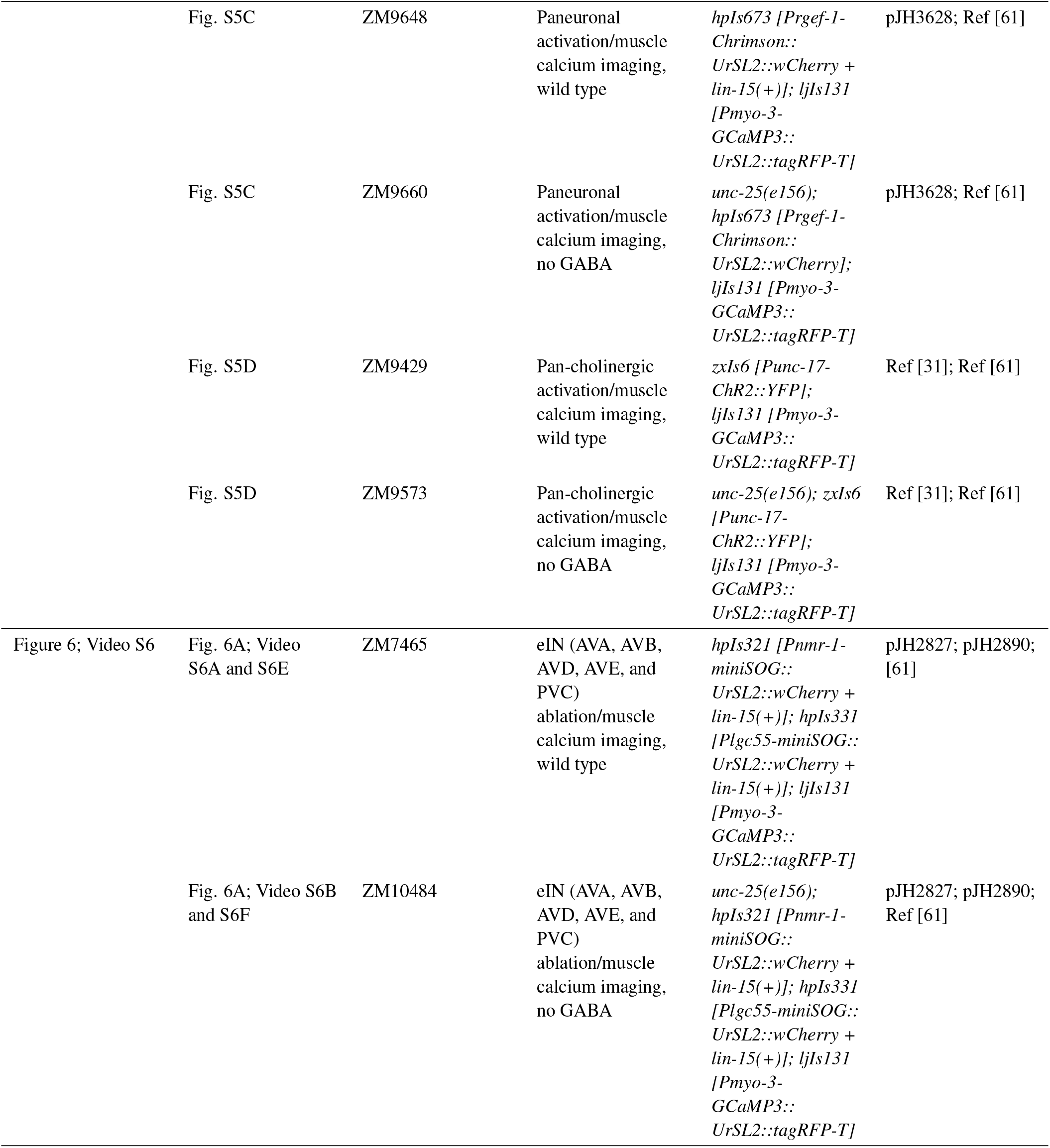
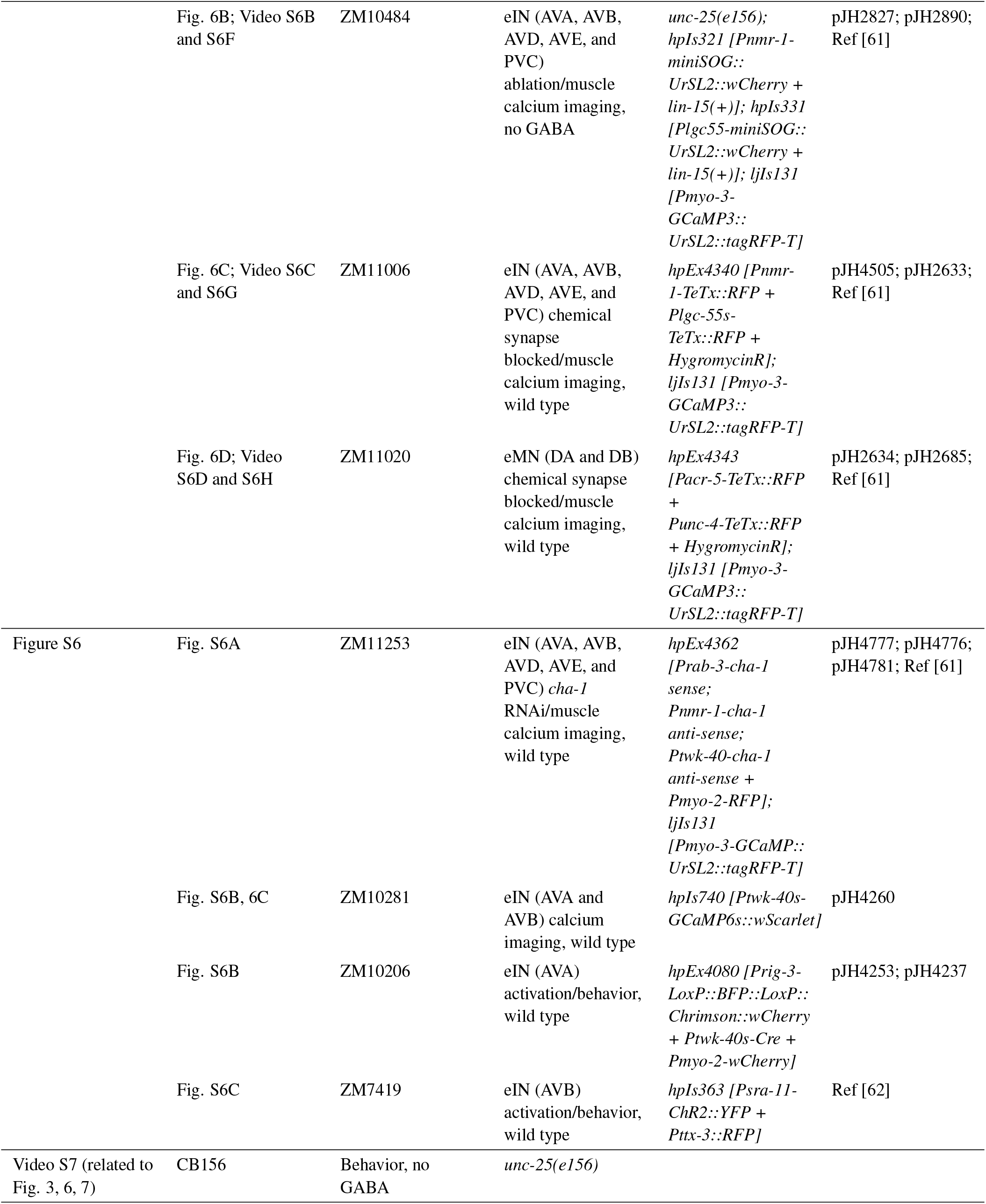
Strains and constructs listed by Figure panels and Videos

### Synchronization of young L1 larvae for all behavioral and imaging experiments

L1 larvae used for behavioral and imaging experiments were collected as follows. On the day of experiment, up to one hundred 3-fold stage eggs from plates with well-fed gravid adults were transferred to a new plate seeded with OP50. Eggs were allowed to hatch for 1hr at 22.5°C and unhatched eggs were removed. Most experiments were carried out 5-6 hours afterwards to ensure that they had not integrated postembryonically derived motor neurons^24^. Some transgenic lines grew a bit slower so experiments were carried out 8-10hr afterwards.

### Electron microscopy of an early L1 stage larva (Figure 1B and S1)

Serial electron microscopy was performed as previously described^63,64^. The L1 larva reconstructed here was at an early L1 stage (∼1-5hr), prior to the beginning of post-embryonic neurogenesis that starts at 6hr. It covers over half of the body,and was manually serial sectioned at 50nm thickness and imaged by TEM at 0.7nm/pixel. Images were stitched into 3D volumes using TrakEM2 Fiji plugin^65,66^, and annotated after skeleton tracing in the CATMAID^67^. A full reconstruction of the ventral and dorsal nerve cords of an newborn L1 larva (1hr) revealed the same pattern and is described in another study^25^.

### Imaging of GABA receptors on L1 larvae (Figure 3C)

GABA receptors were visualized by either direct imaging of tagged fluorescent proteins or immunostaining. For GABA_A_ receptor UNC-49, UNC-49 is tagged with RFP and visualized for RFP fluorescence. The larvae were fixed with 2% paraformaldehyde and 25% methanol to mitigate autofluorescence. Immunostaining of L1 larvae for GBB-1::GFP and GBB-2::GFP with anti-GFP antibodies after fixation was carried out according to [68].

### Analyses of swimming animals (Figure 1C)

Light-field recording of swimming wild type L1 larvae was performed in a PDMS miniature well. A droplet of M9 was mounted onto the center of the well or chamber, and L1 larvae were individually transferred into the droplet. With a cover-slip mounted, the PDMS well was placed on a glass slide and recorded with a 4x objective with a customized compound microscope^19^ for 68s at 26Hz. After images were acquired, body postures were manually scored frame-by-frame. Ventral side of the larva was identified by the presence of anus and/or the gonad. A dorsal-biased bending posture was defined by a convex towards the dorsal side from the neck (30%) to the tail. A ventral-biased bending posture was defined by a convex towards the ventral side from the neck (30%) to the tail. Other postures include both dorsal and ventral bending and were deemed ‘unbiased’. Frames where neither the anus nor the gonad could be confidently identified were discarded. The total number of frames each larva spent in one of three postures was normalized against the total number of frames for each recording.

### Behavioral imaging of swimming L1 larvae (Videos S1A and S7B)

Example movies for swimming wild type and *unc-25* L1 larvae were recorded in a chamber made with a thin circle of vaseline between two glass slides. The chamber was filled with S-basal culture media and multiple L1 larvae. Recordings were carried out with a 10X objective of a Zeiss dissecting microscope (V16) and a Basler aca2500-60um CCD camera with the Pylon viewer program, for 2-5min at 10Hz.

### Calcium imaging of muscle and curvature analyses of crawling animals (Figures 1D, 1E, and 3F; Videos S1C and S3C)

L1 Larvae expressing GCaMP3 and tagRFP separately in body wall muscles were used in these experiments. Dual-color recording of GCaMP3 and mCherry was performed using an in-house ImageJ plugin as described^19,69^. L1 larvae were mounted with a small droplet of M9 or halocarbon oil. Recordings were performed with 20x or 40x objectives, at 26Hz, 20Hz or 10Hz, for 68s, 90s or 180s, respectively, typically by manual stage tracking to keep moving animals in the center of view.

Post-imaging analyses used a customized MATLAB script. The head and tail are manually assigned in the first frame of the recording and tracked afterwards. The contour of the worm is extracted from images. Its mid-line was divided into 100 segments (0 for the head and 100 for the tail), along the anterior-posterior axis, and further segmented into the dorsal and ventral sub-segments across the mid-line. Averaged fluorescent intensity within each segment is used as proxy for the activity for this muscle segment. Bending curvatures were derived from the derivatives of the tangent angles at each segment.

### Calcium imaging of neurons and curvature analyses in crawling animals (Figures 2A, 2B, 3A, 3B, S6B, and S6C; Videos S2 and S3A)

L1 Larvae expressing GCaMP::RFP fusion in neurons were used in these experiments. Recordings were acquired on a customized LED-powered wide-field tracking compound microscope with L1 larvae slowly crawling on 2% agarose pads under a coverslip. Dual-color recording of GCaMP6 and mCherry was performed using an in-house ImageJ plugin as described^19,69^. L1 larvae were mounted with either a small droplet of M9 or halocarbon oil. Recordings were performed with 40x or 63x objectives, at 26Hz, 20Hz or 10Hz, for 68s, 90s or 180s, typically by manual stage tracking.

Calcium imaging analyses was carried out using a MATLAB script modified from [14] that tracks individual regions of interest (ROI). Intensity of individual ROI in the green and red channels was used to calculate calcium dynamics. Curvature at individual ROI was estimated by angles between its immediate neighbours.

### Calculation of phase differences between calcium signals and bending (Figures 1E, 3F, and S2)

Phase differences between dorsal muscle activity, ventral muscle activity, and curvature were carried out using the Hilbert transform, through the MATLAB’s hilbert function. Time-series were converted to a complex-number representation, which was then used to estimate the phase angles at each time-point. Distribution of the differences between phase angles from all samples was then plotted as polar histograms.

### A simple strategy for simultaneous optogenetic stimulation and calcium imaging (Figures 2C, 2D, 3C, 3D, 3E, 4A, 4B, 4D, 4E, 4F, S4, 5, and S5; Videos S3B, S4, and S5)

Simultaneous functional imaging and optogenetic stimulation is technically challenging due to the overlapping excitation spectra for opsins and genetic calcium sensors. Two strategies to address this issue are to spatially separate the illumination^70,71^ or to attempt spectral separation of opsins and calcium sensors^71–76^.

We approached simultaneous optogenetic stimulation and calcium imaging with a simpler goal. Because calcium readout exhibits significant latency compared to the light response of opsin channels, and opsins can be activated by the same illumination spectra for GCaMP and RFP, the initial change of calcium signals should already reflect the cell’s response to opsin stimulation. We validated this strategy with a characterized connection (cholinergic motor neurons to their downstream GABAergic motor neurons in adults), and then applied it to examine the complete L1 motor circuit.

The following protocol conducts bi-directional (activation and inhibition) functional manipulation with any combination between all tested opsins (ChR2, Chrimson, Halorodopsin, GtACR2) and calcium sensors (GCaMP::RFP and chameleon). It can be carried out on wide-field fluorescent as well as confocal microscopes. The following protocol is not meant to probe dynamics of circuit connectivity. It is a simple but effective assay to establish causality and functional connectivity between neurons.

We used the same wide-field fluorescent microscope for calcium imaging. Strains that express opsins in upstream cells and calcium sensors in downstream cells were cultured on all-trans retinal (ATR)-supplemented plates for multiple generations. A control group was cultured on plates without ATR. For experimental groups, particularly strains generated from the Cre-loxP system, animals were cultured on plates supplied with higher concentration ATR (250*μ*M) to increase the strength of optogenetic stimulation. On the day of experiments, L1 larvae were mounted on slides as described for neuron calcium imaging, except that they were immobilized by limited M9 solution. One minute after samples were mounted, recording was carried out initially without the LED light. During the recording, the LED light (serving both for calcium imaging and opsin stimulation) was turned on and off, with an on phase longer than 3 seconds, and off phase ranging from 5-20s. Sampling rates ranged from 10-26Hz. Calcium imaging analyses were carried out as described above. Only the first 3-6s recording for each epoch was used for analyses.

### Calcium imaging and curvature analyses upon eIN and eMN manipulation (Figures S3 and 6; Videos S6A and S6B)

#### Calcium imaging of freely-crawling animals (Figure 6; Videos S6A and S6B)

Calcium imaging recordings for Figure 6 were performed with unconstrained L1 larvae, hatched on a thin lawn of OP50 of 2% agarose plates with S-basal buffer. Images were acquired with a Nikon spinning disk Confocal microscope operating in a wide-field mode with a 10x objective at 10fps. The setup was equipped with a dual view system and Andor Zyla VSC-07720 camera. Toptica’s multi-laser system was used for illumination at 10mW power. Imaging analyses was carried out using a MATLAB script as described in the above section to acquire dorsal and ventral muscle calcium signals and curvature.

#### Photo-ablation of neurons by miniSOG (Figures S3, 6A, and 6B; Videos S6A and S6B)

Transgenic animals that express blue-light activated miniSOG protein were used in these experiments, performed and verified as described in [13].

Briefly, to visually examine ablation efficiency, miniSOG was co-expressed with SL2-RFP to tag the to-be-ablated neurons and neurites. Cells that activated apoptosis usually appear vacuolar after the light treatment. But the complete disappearance of soma and neurites could only be observed towards the end of the larval stage.

A home-made chamber delivers the blue LED light, dissipates heat, and supplies oxygen by ventilation during illumination. For each experiment, 1hr post-hatching larvae on a seeded plate were placed in the chamber upright with its lid removed. The optimal exposure time was experimentally determined for each strain to ablate cells without arresting larva growth. The eIN ablation strain required 30min exposure. After illumination, animals were allowed to recover in darkness for 4hr before behavioral and calcium imaging experiments. The control group larvae were similarly hatched on a seeded plate and maintained alongside the experimental group (larvae plates) after their LED treatment. The larger variation in these control groups (compared to the Control groups in Figures 6C and 6D, which were wildtype animals) was likely caused by their exposure to blue and red laser during calcium imaging experiments.

It takes up to two days for ablated neurons to anatomically disintegrate and absorbed by other tissues. To verify successful ablation of neurons, after each larva was imaged, it was recovered and transferred to individual NGM culture plates with OP50 to grow for two days into adulthood. They were then examined for RFP signals using a compound microscope. Only data recorded from those with absence of RFP in targeted neurons were used for analyses.

### Wide-field imaging and analyses of crawling animals (Figures S6B and S6C; Videos S1B, S6E, S6F, S6G, S6H, and S7A)

*On plate recording of free crawling L1s (Videos S1B, S6E, S6F, S6G, S6H, and S7A)*. Recordings were performed using a customized wide-field, white-light compound microscope^19^, one larva at a time, with a thin-layer of OP50 bacteria food on a NGM plate. Larva movements manually tracked to stay in the center of view during recording, typically for 75sec at 26Hz.

*On plate recording free crawling L1s upon optogenetic stimulation (Figures S6B and S6C)*. For each set of optogenetic experiments, the control and experimental group contained animals of the same genotype, with the control group cultured on plates without all-trans retinal (ATR)^77^. All larvae were recorded using image plates that were unseeded with OP50 and without ATR, typically for 75s at 26Hz.

Animals in the experimental group was cultured on NGM plates supplied with 0.1mg/ml ATR for at least two generations. Individual Ll larvae were transferred to and recorded using the same unseeded imaging plate as described above. L1 larvae from the control group was recorded on a separate imaging plate.

All animals were subjected to stimulation from an LED light source controlled by a customized ImageJ plugin^19^. The stimulation protocol was 10s light-off (pre-stimulation), 4 consecutive epochs of stimulation consisting of 10s light-on and 5s light-off, followed by 10s light-off (post-stimulation).

*Velocity measurements of crawling animals(Figures S6B and S6C)*. Velocity was carried out using in-house MATLAB scripts https://github.com/zhen-lab/Beta_Function_Analysis. Briefly, contours of recorded L1 larva images above were segmented and divided into 100 segments along the anterior-posterior axis. Orientation (head, tail, ventral, and dorsal) were manually assigned in the first frame of the recording and tracked automatically. Errors were fixed by manual curation afterwards. Instantaneous velocity was measured by the displacement of centroid between frames; those towards the head are forward movement and those towards the tail are backward movement.

### A computational model (Figure 7B)

We modeled each segment of the L1 motor circuit using the equations below (1-5), which is schematized in Figure 7B. The model is inspired by the Morris-Lecar model of the Barnacle giant muscle fiber^78^. It’s a reduced order nonlinear model with an instantaneous calcium current, a potassium current with an activation variable, and a passive leak current. A similar model was used previously to study adult *C. elegans* excitatory motor neurons^16^. We model 6 segments coupled to each other proprioceptively, corresponding to the 6 DD NMJs. *V*_*di*_, *M*_*di*_, *M*_*vi*_ are the membrane potentials for the eMN, dorsal muscle and ventral muscle in segment *i* respectively. *n*_*i*_ is the activation variable for the slow potassium current and *κ*_*i*_ is the segment’s curvature. We implicitly model the activity of DD iMNs via an inhibitory synapse on ventral muscles by eMNs. We did not include the extrasynaptic inhibition to dorsal muscles in our model, because it was not necessary for the generation of a symmetric oscillatory gait. This also recapitulates our experimental findings, where inhibitory feedback on to dorsal muscles is modulatory and *gbb-1* mutant larvae did not exhibit overt motor defects.

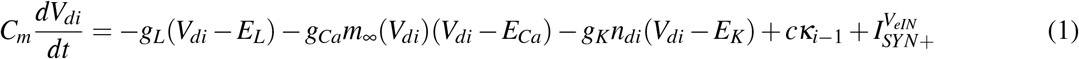

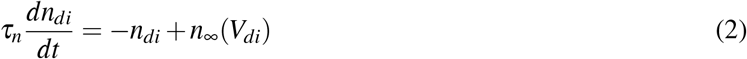

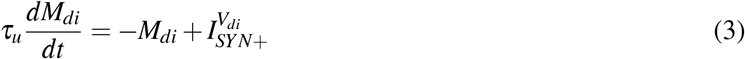

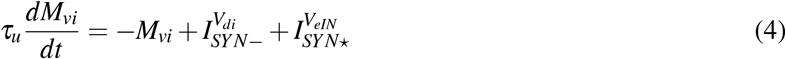

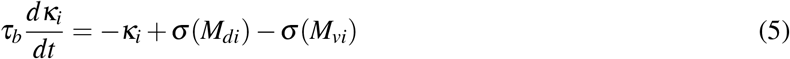

Where, *E*_*L*_ = −60 mV, *E*_*Ca*_ = 60 mV, *E*_*K*_ = −70 mV are the leak, calcium and potassium reversal potentials respectively. Terms *g*_*L*_ = 100 pS, *g*_*Ca*_ = 400 pS, *g*_*K*_ = 500 pS are the maximal leak, calcium and potassium conductances and *C*_*m*_ = 3 pF is the membrane capacitance. Timescale parameters were set as *τ*_*n*_ = 30 ms, *τ*_*u*_ = 85 ms, *τ*_*b*_ = 10 ms The function 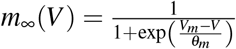 with *θ*_*m*_ = 10.25 mV,*V*_*m*_ = −29 mV. And 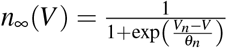 with *θ*_*n*_ = 20 mV,*V*_*n*_ = −55 mV. While, *σ* (*V*) = *α*(tanh(*β* ∗ *V*) + 1) with and *α* = 1000 mm^−1^ and *β* = 0.01 mV^−1^ converts muscle activities to curvature.

The terms 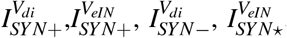, respectively are the synaptic currents from the excitatory synapse from eMN to dorsal muscles, excitatory synapse from eINs to the eMN, inhibitory synapse from eMNs to ventral muscles, and the excitatory extrasynaptic input from eIN to ventral muscles. The synaptic currents are based on the graded synapse model used in^79^, and are defined as follows.

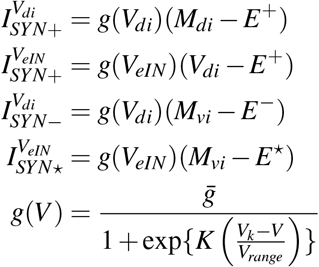

With 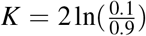, set such that the conductance varied from 10% to 90% of its maximum value over the presynaptic voltage range set by *V*_*range*_. Other parameters were set as 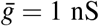,*V*_*k*_ = −30 mV,*V*_*range*_ = 20 mV. Terms *E*^+^, *E*^−^, *E*^*⋆*^ are the reversal potentials for the excitatory receptor, inhibitory receptor and the extrasynaptic receptor. We set *E*^+^ = −10 mV. *E*^−^ and *E*^*⋆*^ were treated as variables and optimized to result in an unbiased gait, such that the mean value of the bending curvature ⟨*κ*_*i*_⟩ is zero. The optimized values were found to be *E*^−^ = −67 mV, *E*^*⋆*^ = 33.5 mV.

## Acknowledgments

We thank Taizo Kawano, Ying Wang, Yan Li, Jin Meng, Sway Chen, Yi Li, Bin Yu, Thomas Sun and Lu Hang for technical support; Albert Cardona for guidance on CATMAID; Michael Koelle and lab members of Mei Zhen, Shangbang Gao and Aravinthan DT Samuel for discussions. This work was supported by the International Human Frontier Science Program Organization RGP0051/2014 (MZ; ADTS), the Canadian Institutes of Health Research Foundation Scheme 154274 (MZ), and the National Institute of Health R35 GM134970 (ADC) and R01 NS093588 (ADC).

## Contributions

MZ and QW conceived the study; MZ supervised the study; YL, TA, AG, BM, WH, JM and MZ designed and performed experiments and analyzed data; DW, BM, DH, ADC, MZ performed EM reconstruction; AC guided annotation; TA developed model; YL, TA, DW and QW developed scripts; YL, TA, ADTS and MZ wrote the paper; all authors read or edited the paper.

## Interests

Authors declare no competing interests.

## Supplemental Information

**Figure S1.**
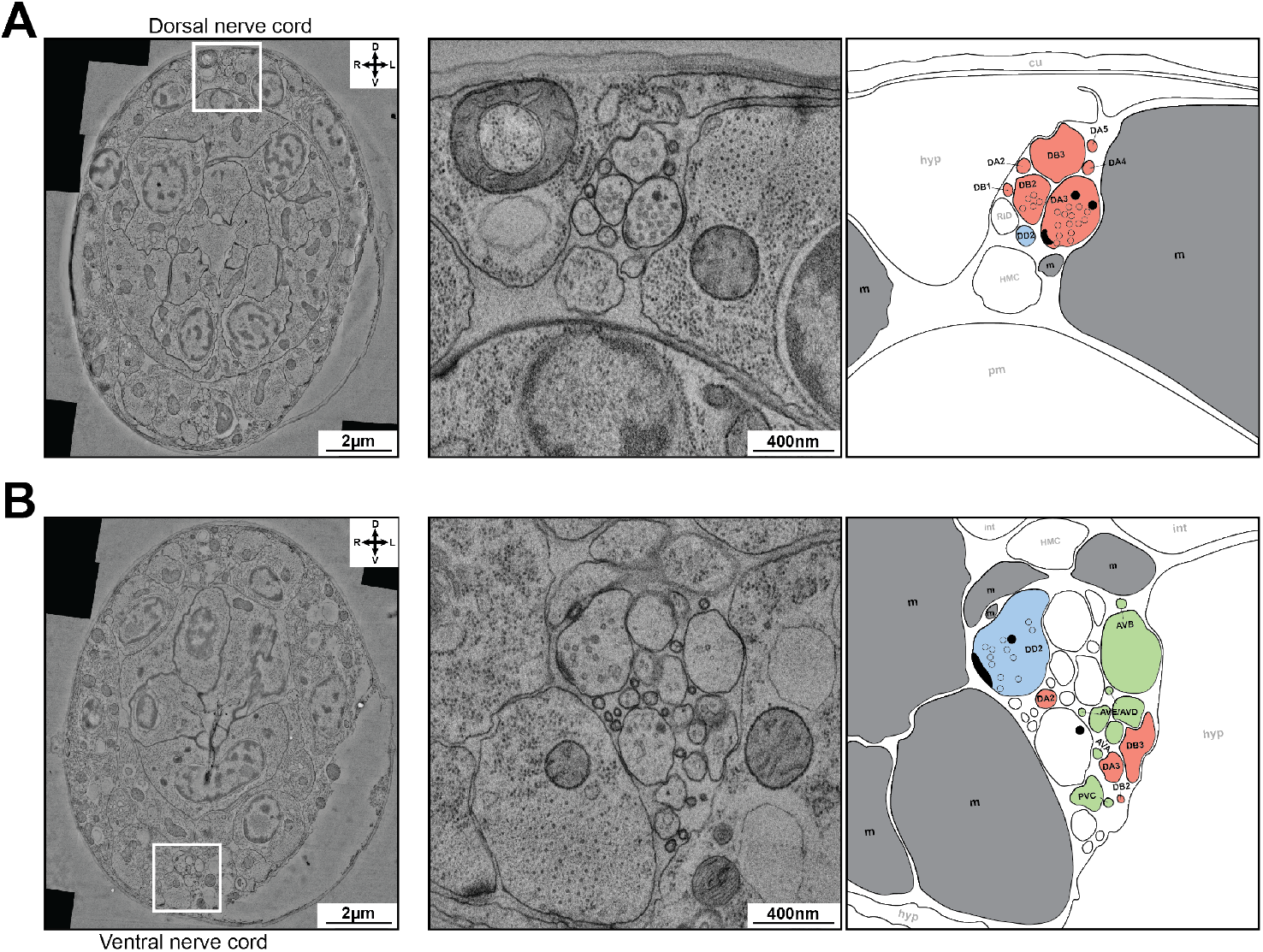
Electron microscopy reconstruction of the dorsal and ventral nerve cord of an 5-6hr L1 larva. (A) Example image of the dorsal nerve cord. (Left) An electron micrograph of the cross-section in transverse plane. The dorsal nerve cord is marked. (Middle) An enlarged view of the dorsal nerve cord. (Right) Cross-sections of individual neurites of motor neurons and muscles of the L1 motor circuit were color-coded and annotated. Red: eMNs; blue:iMNs; grey: muscles. One eMN (DA3) makes a dyadic NMJ to the muscle and iMN (DD2), with an active zone (black crescent), a cluster of synaptic vesicles (clear circles) and sparse dense core vesicles (dark circles). (B) Example image of the ventral nerve cord, similarly illustrated and annotated as in (A). Neurites for eINs, which are absent from the dorsal nerve cord (A), are color-coded green. An iMN (DD2) makes a NMJ to muscles. In right panels, cross-sections of other neuron (RID) and cell (HMC) of the L1 motor circuit in the dorsal nerve cord (A) and ventral nerve cord (B) were labelled by opaque letters. *Abbreviations:* m: muscle; pm: pharyngeal muscle; hyp: hypodermis; cu: cuticle; int: intestine.

**Figure S2.**
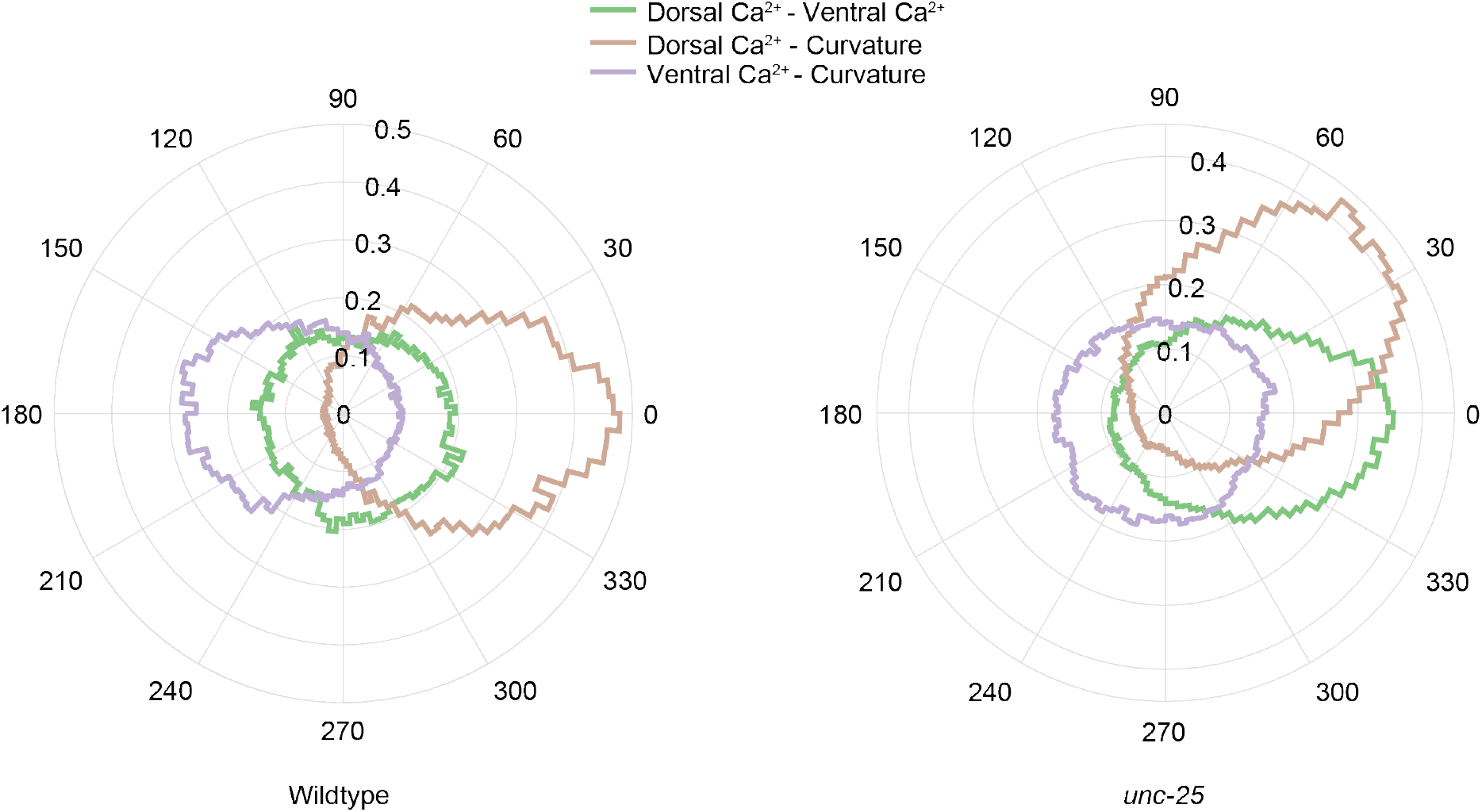
Calcium dynamics of dorsal muscles tracks bend waves in *unc-25* mutant larvae. Polar histograms of the phase differences between curvature, dorsal muscle activity, and ventral muscle activity (segments 33-95) for crawling wildtype and *unc-25* larvae imaged under the same condition. (Left) In wildtype larvae, dorsal muscle activity was in phase with body bending and ventral muscle was anti-phasic with body bending. (Right) In *unc-25* larvae, ventral muscles had no phase relationship with curvature, but dorsal muscle activity tracked body bending with phase lag. N = 10 recordings for each group.

**Figure S3.**
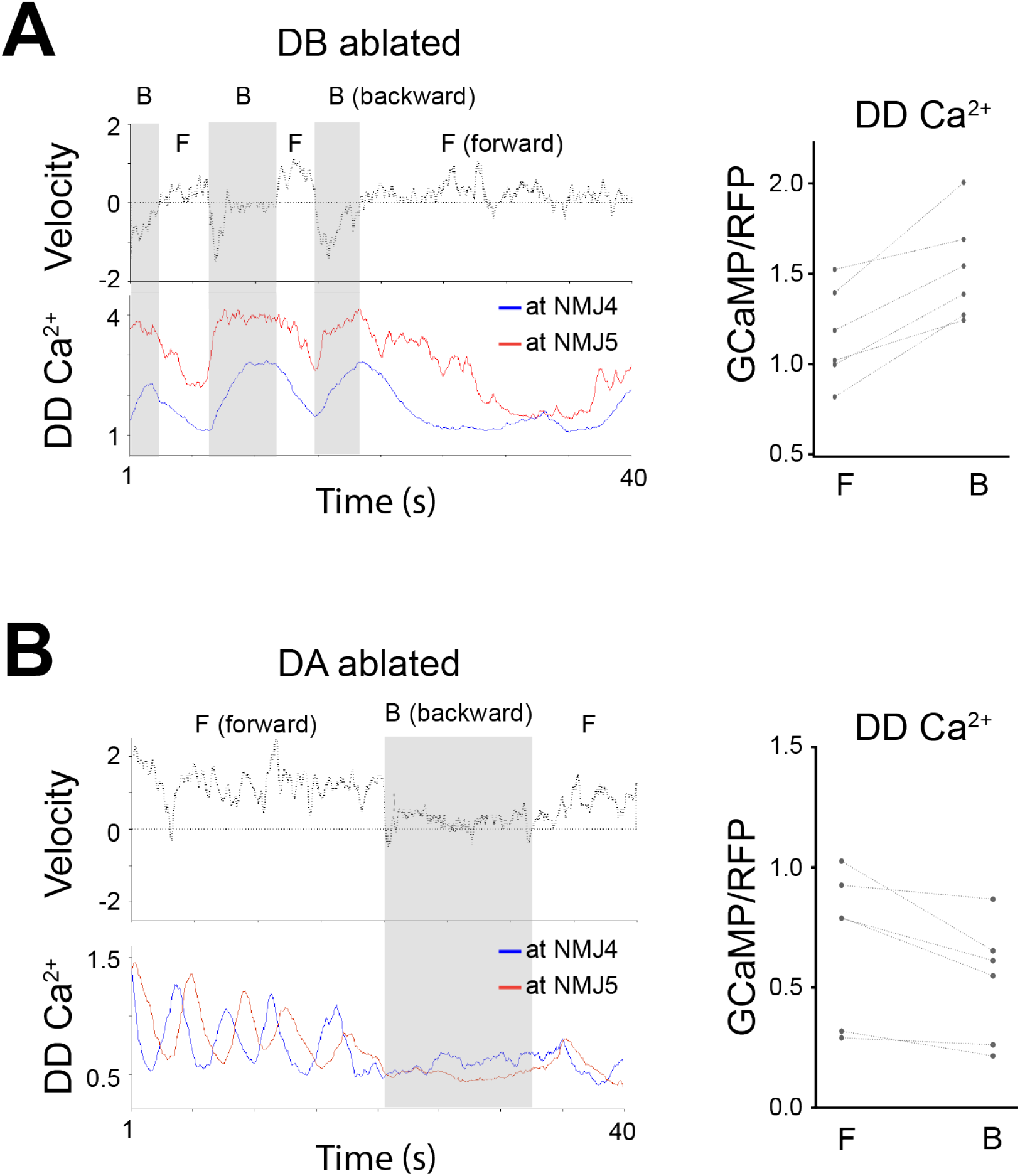
Ablation of eMNs reduces iMN activity during directional movement. (A) (Left) Example of velocity and DD NMJ calcium of an L1 lava upon DA motor neuron ablation. This larva still transited between forward and backward movement, with attempted reversals shown as pauses. (Right) Mean DD activity during forward and backward movement, with lines connecting epochs from the same animals. (B) (Left) Example of velocity and DD NMJ calcium of an L1 lava upon DB motor neuron ablation. This larva still transited between forward and backward movement, with attempted forward movement shown as pauses. (Right) Mean DD activity during forward and backward movement, with lines connecting epochs from the same animals.

**Figure S4.**
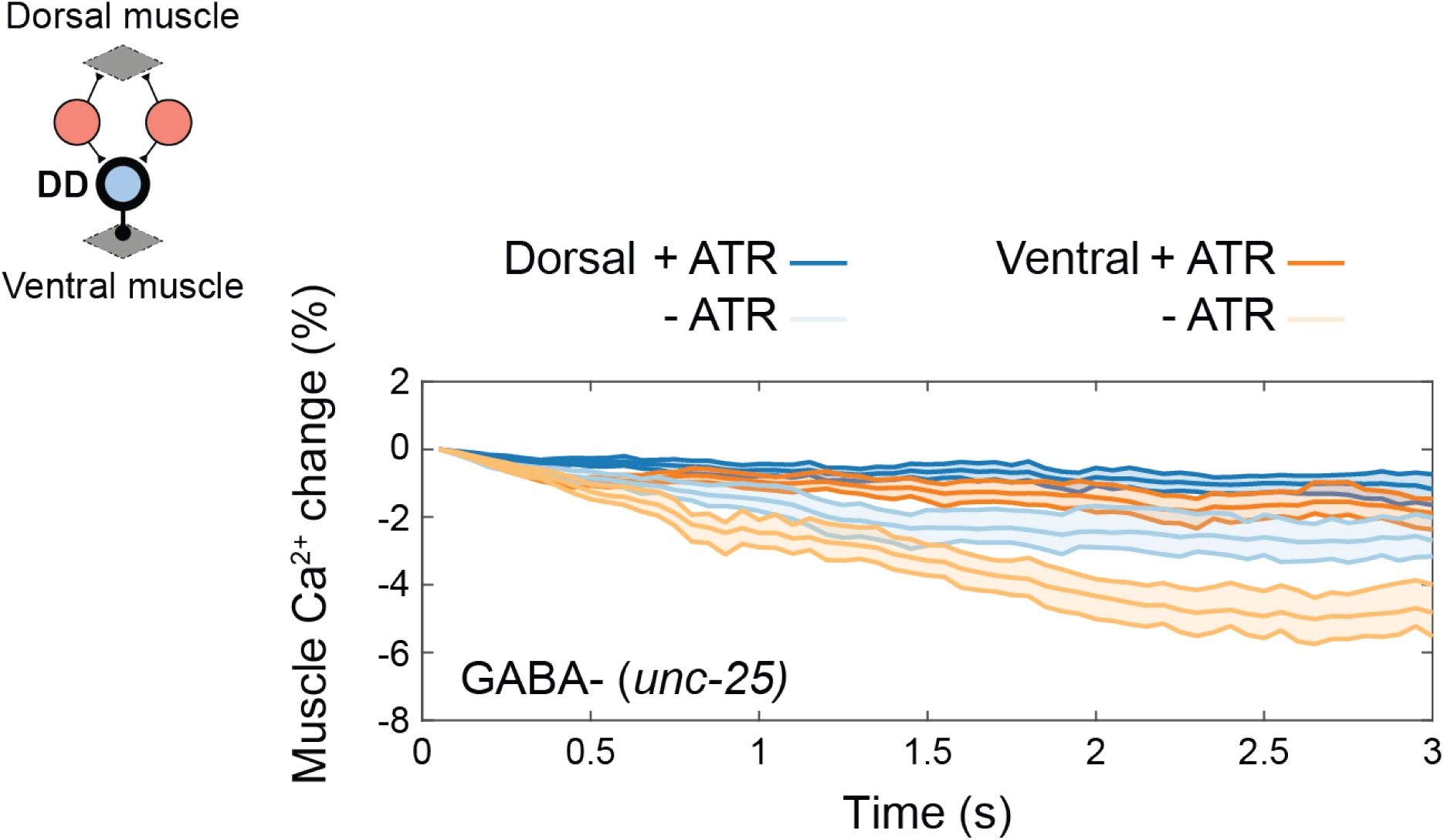
Activation of iMNs does not significantly reduce ventral and dorsal muscle activity in mutant larvae that do not synthesize GABA. (Inset) Schematic of simultaneous optogenetic activation of iMNs and muscle calcium imaging. Y-axis plots percentage changes of the muscle activity from *t* = 0. Control group (-ATR): 34 stimulation epochs from 12 larvae; Experimental group (+ ATR): 15 stimulation epochs from 5 larvae.

**Figure S5.**
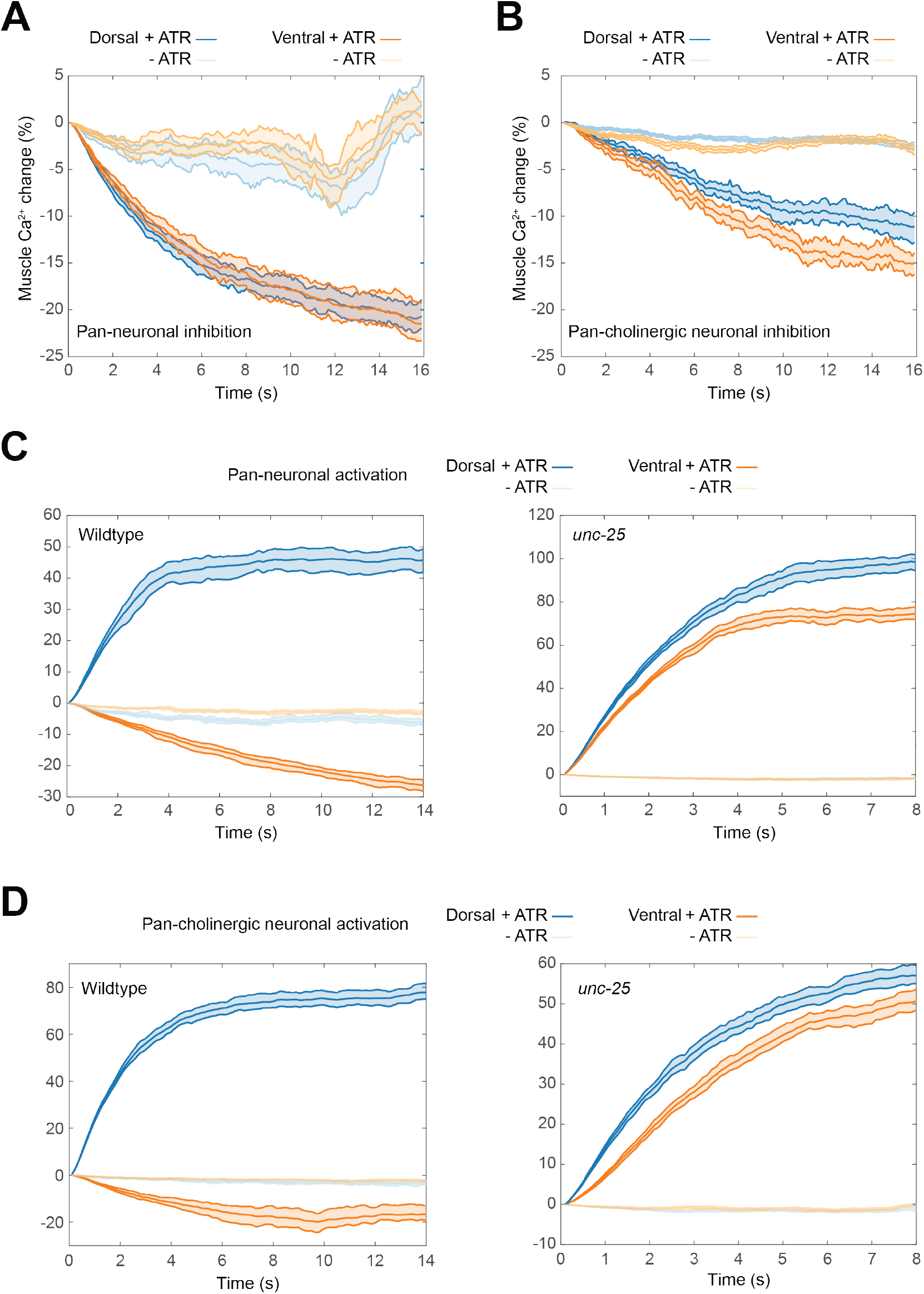
Effect of optogenetic manipulation of groups of neurons on dorsal and ventral muscles. (A-B) Muscle calcium responses upon optogenetic silencing of all neurons (A) or all cholinergic neurons (B) in wildtype larvae. Y-axis plots percentage changes of the muscle activity from *t* = 0. Panneuronal inhibition: the control group (-ATR): 18 stimulation epochs from 6 larvae; the experimental group (+ ATR): 26 stimulation epochs from 12 larvae. Cholinergic neuron inhibition: the control group (-ATR), 16 stimulation epochs from 3 larvae; the experimental group (+ ATR), 27 stimulation epochs from 5 larvae. (C) Muscle calcium responses upon optogenetic activation of all neurons in wildtype (left) or *unc-25* (right) larvae. Wildtype group: - ATR: 17 stimulation epochs from 9 larvae; + ATR: 18 stimulation epochs from 10 larvae. *unc-25* group: - ATR: 14 stimulation epochs from 9 larvae; + ATR: 21 stimulation epochs from 11 larvae. (D) Muscle calcium responses upon activation of all cholinergic neurons in wildtype (left) or *unc-25* (right) larvae. Wildtype group: - ATR, 15 stimulation epochs from 7 larvae; + ATR, 19 stimulation epochs from 10 larvae. *unc-25* group: - ATR, 18 stimulation epochs from 6 larvae; + ATR, 30 stimulation epochs from 10 larvae.

**Figure S6.**
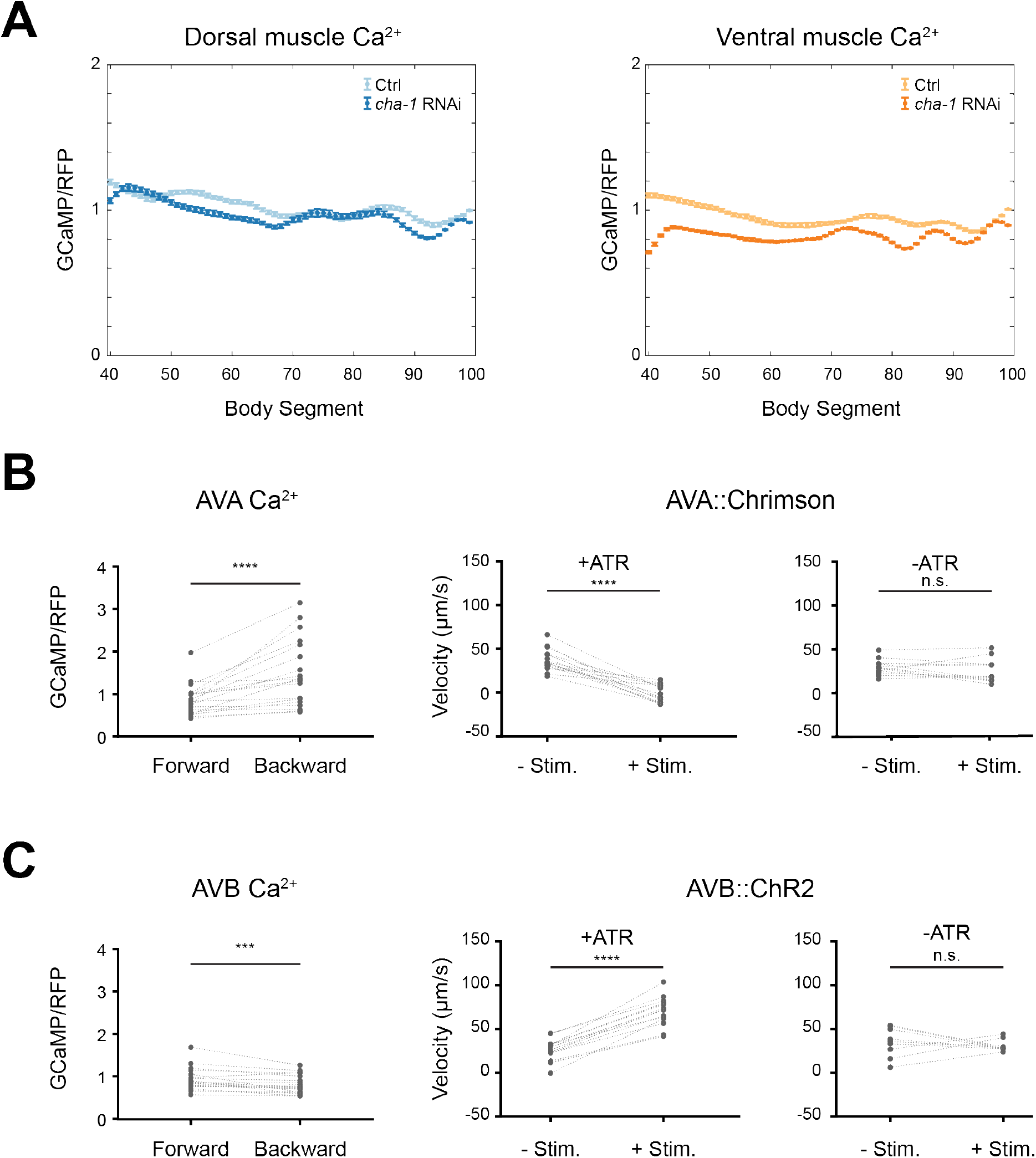
Effect of genetic and optogenetic manipulation of eINs in L1 larvae. (A) RNAi-mediated knockdown of *cha-1* in all eINs leads to stronger reduction of ventral muscle activity. Dorsal (left panel) and ventral (right panel) calcium signals compared between larvae carrying only the calcium reporter (Ctrl) and larvae that carried an additional transgene expressing anti-sense *cha-1* in all eINs (*cha-1* RNAi). In these larvae, segments 40-100 (instead of 33-100, as in Figure 6) constitute the body because they were slightly younger. (B) (Left panel) The AVA eIN exhibited higher activity during backward movement. n = 19 larvae. (Middle/right panels) Optogenetic activation of AVA led to reversals. Mean velocities during non-stimulated and stimulated phases were compared in the control group (-ATR) (n = 12 larvae) and the experimental group (+ ATR) (n = 15 larvae). **** p < 0.0001. (C) (Left panel) The AVB eIN exhibited higher activity during forward movement (n = 19 larvae). *** p = 0.0006. (Middle/right panels) Optogenetic activation of AVB led to increased forward velocity. Control group (-ATR): n = 9 larvae. Experimental group (+ ATR): n = 15 larvae. **** p < 0.0001. All p-values were calculated using the Wilcoxon matched-pairs signed rank test.

## Movie captions

**Video S1 (Figure 1): Movements and muscle calcium dynamics of the L1 larvae**.

(A) (Figure 1C) An L1 larva swimming with alternating body bends in M9 solution. In the first frame, head is up and tail is down. A lump of bacteria food anchored the animal. (B) An L1 larva crawling with on an NGM plate with a thin layer of bacteria food. In the first frame, head is down, tail is up, dorsal is left and ventral is right. (C) (Figures 1D and 1E) Muscle calcium activity of an L1 larva crawling on an agarose pad under the coverslip. *Upper panels* Calcium imaging of a crawling L1 larva that expressed GCaMP3::RFP in boy wall muscles. Head: left, tail: right, dorsal: up, and ventral: down. GCaMP3/RFP ratio was segmented along the body. Color bar and the Y-axis of the lower panel have the same scale. Dashed line denotes the segment that separates the head and body. *Lower panel* Calcium signals in dorsal (blue) and ventral (red) muscles over time. X-axis represents body segments (head: 1, tail: 100); dashed line denotes the segment that separates the head and body. Y-axis represents the GCaMP3/RFP ratio at each segment.

**Video S2 (Figures 2A and 2B): eMN activities increase sequentially during L1 larva’s directional movements**.

(A) (Figure 2A) DA motor neurons increase their activities sequentially during backward locomotion. Head: left; tail: right; dorsal: up; and ventral: down. Left-pointing arrow denotes periods of forward movement; right-point arrow denotes periods of backward movement. GCaMP6 and RFP signals from DA3-DA9 somata were tracked. Grey bars denote periods of backward movement. (B) (Figure 2B) DB motor neurons increase their activities sequentially during forward locomotion. GCaMP6 and RFP signals were present in both eMN classes, but stronger in the DB class. GCaMP6 and RFP signals from DB4-DB6 somata were tracked. Grey bars denote periods of forward movement.

**Video S3 (Figure 3A, 3B, 3D, 3E, 3F, and S2A): GABAergic iMNs relax and couple dorsal and ventral muscle activities**.

(A) (Figures 3A and 3B) NMJs from iMNs are activated during both forward and backward movements. DD motor neurons activities were shown by calcium imaging of a crawling L1 larva expressing GCaMP6::RFP in DD motor neurons. Head: left; tail: right; dorsal: up; ventral: down. Left-pointing arrow denotes periods of forward movement; right-point arrow denotes periods of backward movement. Grey bars denote periods of backward movement. GCaMP6 and RFP signals from NMJs of DD5 and DD6 were tracked. (B) (Figures 3D and 3E) Inhibition of iMNs in wildtype larvae elevates dorsal and ventral muscle activity with different spatial patterns. *Upper panels* Calcium imaging of an immobilized L1 larva that expressed GCaMP6::RFP in muscle cells and Archaerhodopsin in DD motor neurons. GCaMP3/RFP ratio was segmented along the body. Color bar and the Y-axis of the lower panel have the same scale. Dashed line denotes the segment that separates the head and body. *Lower panel* Calcium signals in dorsal (blue) and ventral (red) muscles over time. X-axis represents body segments (head: 1, tail: 100); dashed line denotes the segment that separates the head and body. Y-axis represents the GCaMP3/RFP ratio at each segment. (C) (Figures 3F and S3A) Larvae without GABA exhibit decoupled muscle calcium signals on dorsal and ventral sides. *Upper panels* Calcium imaging of a crawling L1 larva that expressed GCaMP3::RFP in boy wall muscles. *Lower panel* Calcium signals in dorsal (blue) and ventral (red) muscles over time.

**Video S4 (Figure 4): iMNs are innervated by eMNs**.

(A) (Figure 4A) iMNs are activated upon eMNs stimulation. Simultaneous calcium imaging of DD motor neurons and optogenetic stimulation of DA and DB motor neurons with an immobilized L1 larva expressing GCaMP6::RFP in DD motor neurons and Chrimson in DA and DB motor neurons. The larva was fed with OP50 bacteria containing all-trans retinal (ATR). The starting frame is the when calcium imaging and optogenetic manipulation occur simultaneously. Head: left, tail: right, dorsal: up and ventral: down. Right panel shows percentage changes of the muscle activity from *t* = 0. NMJs from DD4 and DD5 were tracked. (B) (Figure 4A) iMNs are not activated upon illumination in the control animal. The same illumination condition in (A) was applied to the larva of the same genotype, but fed with OP50 without ATR. (C) (Figure 4A) iMNs are inactivated upon eMNs silencing. Simultaneous calcium imaging of DD motor neurons and optogenetic silencing of DA and DB motor neurons with an immobilized L1 larva expressing GCaMP6::RFP in DD motor neurons and Archaerhodopsin in DA and DB motor neurons. The larva was fed with OP50 containing ATR. NMJs from DD5, DD6 and DD7 were tracked. (D) (Figure 4A) iMNs are not inactivated upon illumination in the control animal. The same illumination condition in (C) was applied to the larva of the same genotype, but fed with OP50 without ATR.

**Video S5 (Figures 5A and 5B): Activation and inactivation of eINs induce changes that differ between dorsal and ventral muscles**.

(A) (Figure 5A) Silencing eINs reduces both dorsal and ventral muscle activities. *Upper panels* Calcium imaging of an immobilized L1 larva that expressed GCaMP3::RFP in body wall muscles and Archaerhodopsin in multiple eINs. *Lower panel* Calcium signals in dorsal (blue) and ventral (red) muscles over time. X-axis represents body segments (head: 1; tail: 100); the vertical dashed line denotes the segment that separates the head and body. Y-axis represents GCaMP3/RFP value at each segment. (B) (Figure 5B) Activation of eINs induces spatially distinct calcium increase in *unc-25* mutant larva’s dorsal and ventral muscles. *Upper panels* Calcium imaging of an immobilized *unc-25* mutant larva that expressed GCaMP3::RFP in body wall muscles and ChR2 in AVA. *Lower panel* Calcium signals in dorsal (blue) and ventral (red) muscles over time.

**Video S6 (Figures 6A, 6B, 6C, 6D): Extrasynaptic transmission from eINs is required for ventral bending**.

(A) (Figure 6A) Reduced ventral muscle activity after ablation of eINs in wildtype larva. *Upper panels* Calcium imaging of an unconstrained L1 wildtype larva that expressed GCaMP3::RFP in body wall muscles after photo-ablation of all eINs by miniSOG. It adopted a dorsally biased posture. Color bar and Y-axis of the lower panel represent the same scale. Dashed line denotes the body segment that separates the head and the body. *Lower panel* Calcium signals in dorsal (blue) and ventral (red) muscles over time. X-axis represents body segments (head: 1; tail: 100); the vertical dashed line denotes the segment that separates the head and body. Y-axis represents GCaMP3/RFP value at each segment. (B) (Figure 6B) Reduced ventral muscle activity after ablation of eINs in *unc-25* mutant larva. *Upper panels* Calcium imaging of an unconstrained L1 *unc-25* larva that expressed GCaMP3::RFP in body wall muscles after photo-ablation of all eINs by miniSOG. It adopted a dorsally biased posture. *Lower panel* Calcium signals in dorsal (blue) and ventral (red) muscles over time. (C) (Figure 6C) Blocking synaptic vesicle release from eINs reduces calcium signals in ventral muscles of the larva. *Upper panels* Calcium imaging of an unconstrained L1 larva that expressed GCaMP3::RFP in body wall muscles and TeTx in all eINs. It crawled with a dorsally biased posture. *Lower panel* Calcium signals in dorsal (blue) and ventral (red) muscles over time. (D) (Figure 6D): Blocking synaptic vesicle release from eMNs reveals uniformed calcium signals in ventral muscles of the larva. *Upper panels* Calcium imaging of an unconstrained L1 larva that expressed GCaMP3::RFP in body wall muscles and TeTx in eMNs. *Lower panel* Calcium signals in dorsal (blue) and ventral (red) muscles over time. (E) Behavior of eIN-ablated wildtype L1 larvae. Multiple wildtype L1 larvae lying on agar plate with curled body postures after photo-ablation of all eINs by miniSOG. They adopted a dorsally biased posture. (F) Behavior of eIN-ablated *unc-25* mutant L1 larvae. Multiple *unc-25* L1 larvae lying on agar plate with curled body postures after photo-ablation of all eINs by miniSOG. They adopted a dorsally biased posture. (G) Behavior of eIN synaptic output-blocked L1 larvae. It crawled with a dorsally biased posture. (H) Behavior of eMN synaptic output-blocked L1 larvae. It adopted a ventrally biased posture.

**Video S7 (Figure 7): Complex bending defects in crawling for the *unc-25* mutant larvae**.

*unc-25* mutant larvae exhibit bending defects that are difficult to quantify for ventral or dorsal bias. Qualitatively, they rest in a body posture with ventral bending across the body. When they are prompted to move, their forward movement does not exhibit overt dorsal or ventral bias, whereas their reversals include rapid ventral coils. Same trends are observed for both crawling (A) and swimming (B).

